# Integrative spatial profiling of 3D genome organization and gene expression in tissue

**DOI:** 10.64898/2026.07.28.741242

**Authors:** Pengfei Guo, Yan Cui, Jincan He, Abraham J. Waldman, Jiaxin Zhu, Yufan Chen, Zhi Huang, Jingtian Zhou, Jennifer E. Phillips-Cremins, Yanxiang Deng

## Abstract

The interplay between 3D genome architecture and transcriptional activity is fundamental to gene regulation. However, existing methodologies cannot simultaneously measure these modalities within intact tissues, limiting our understanding of how genome organization coordinates transcriptional programs across diverse cell types and spatial microenvironments. Here, we introduce Spatial Hi-C-RNA, a spatial multi-omics technology that enables the genome-wide co-mapping of chromatin conformation and transcriptome directly from the same tissue section at near-single-cell resolution. Applied to the mouse embryo and adult brains, Spatial Hi-C-RNA generated high-resolution tissue maps revealing that chromatin organization and gene expression jointly define spatially coherent domains aligned with histological structures. While concordant features were observed across modalities, distinct domain patterns also emerged, indicating that chromatin structure and transcription each contribute complementary layers of spatial regulation. We further demonstrated the robustness and biological insight of Spatial Hi-C-RNA in human melanoma, where both modalities delineated tumor boundaries and microenvironmental niches. Notably, chromatin maps revealed fine-scale tumor subdomains undetectable by transcriptomic profiling alone, highlighting the added resolution provided by spatial chromatin architecture. Integrated analysis revealed that multiscale 3D genome features, from A/B compartments and topologically associating domains to chromatin loops, are closely coupled with domain- and cell-type-specific transcriptional programs. In addition, Spatial Hi-C-RNA resolves spatiotemporal dynamics underlying embryonic lineage specification and tumor progression. Together, these capabilities extend the spatial omics landscape beyond transcriptome and epigenome profiling to the level of chromatin organization, establishing an integrative framework for understanding tissue biology across development and disease.

## INTRODUCTION

The three-dimensional (3D) organization of the genome is a fundamental layer of gene regulation.^1,2^ The spatial folding of the genome brings distal regulatory elements into proximity, enabling the precise coordination of transcriptional programs that underpin cellular identity and states.^3–5^ This hierarchical architecture—spanning chromosome territories^6^, A/B compartments^7^, topologically associating domains (TADs)^8,9^ and chromatin loops^10^—is essential for development and differentiation, while its disruption drives pathologies such as cancer and neurodevelopmental disorders.^11–13^ Therefore, to fully decipher how genome structure governs gene expression, it is essential to measure chromatin architecture and transcriptional output simultaneously within the same cells.

Recent advances in single-cell Hi-C have provided unprecedented insights into the cell-to-cell heterogeneity of the 3D genome organization.^14–17^ Technologies capable of co-profiling chromatin contacts and RNA have begun to link chromatin folding to transcription, revealing key principles of enhancer–promoter communication and architectural heterogeneity.^18–22^ However, these approaches inherently require tissue dissociation, leading to the loss of spatial contexts that are essential for understanding genome regulation *in vivo*.

Emerging spatial omics technologies now enable the *in situ* profiling of DNA methylation, chromatin accessibility, histone modifications, transcripts, and proteins while preserving tissue architecture.^23–31^ These methods have transformed our understanding of developmental patterning, brain organization, and tumor ecosystems.^32–34^ Furthermore, spatial context refines cell-type annotation by integrating positional information with molecular states, allowing for the discrimination of closely related subtypes that exhibit overlapping transcriptional or epigenetic profiles.^25,35,36^ While imaging-based approaches have achieved simultaneous detection of specific 3D genomic loci and RNA transcripts,^37,38^ they remain inherently targeted. These methods are restricted to pre-selected genomic regions and typically lack the ability to capture unbiased, genome-wide chromatin folding patterns. Consequently, a scalable technology capable of measuring genome-wide chromatin conformation and the whole transcriptome simultaneously in intact tissues is still lacking. Without a comprehensive spatial view of 3D genome folding, how physical genome organization orchestrates transcriptional programs within their native spatial context remains elusive.

Here, we introduce Spatial Hi-C-RNA, a spatial multi-omics technology that simultaneously maps 3D chromatin architecture and whole-transcriptome profiles from the same tissue section at near-single-cell resolution. By integrating Hi-C with spatially resolved RNA sequencing, Spatial Hi-C-RNA captures multiscale genome organization alongside corresponding transcriptional states in their native tissue contexts. Applied to the developing mouse embryo, adult mouse brains, and human melanoma, this platform reveals tissue-specific spatial architecture in which chromatin organization and gene expression jointly define spatial domains, as well as complementary modality-specific patterns that highlight distinct layers of genome regulation. In melanoma, chromatin maps reveal fine-scale malignant subdomains that are not detectable by transcriptomic profiling alone, demonstrating the added resolution provided by spatial genome architecture. Together, Spatial Hi-C-RNA provides a comprehensive framework for linking 3D genome architecture to transcriptional regulation and cellular function within intact tissues, illuminating the spatial and temporal principles that guide gene regulation across development and disease.

## RESULTS

### Development and benchmarking of Spatial Hi-C-RNA for joint profiling of chromatin architecture and gene expression in tissues

Spatial Hi-C-RNA integrates deterministic microfluidic *in situ* barcoding,^39^ high-throughput chromatin conformation capture,^19,20,40^ and RNA sequencing to simultaneously profile 3D genome organization and gene expression within intact tissue sections (Figure 1A). Briefly, a fresh-frozen tissue section was fixed using a dual-crosslinking strategy^41^ (formaldehyde and disuccinimidyl glutamate) to preserve chromatin structure. Following permeabilization, reverse transcription was performed using a biotinylated poly(T) primer containing a unique molecular identifier (UMI) and a universal ligation linker, synthesizing complementary DNA (cDNA) from captured messenger RNA (mRNA) *in situ*. Crosslinked chromatin was subsequently fragmented using a cocktail of three restriction enzymes (DpnII, DdeI, and HinfI) and subjected to proximity ligation. To minimize open chromatin bias and enhance Tn5 tagmentation efficiency, the tissue section was treated with sodium dodecyl sulfate (SDS) prior to inserting the universal ligation linker into genomic DNA via Tn5 transposition.^20^ Spatially resolved barcoding was achieved by sequentially applying two orthogonally oriented polydimethylsiloxane (PDMS) microfluidic chips to the tissue section. Through two rounds of ligation, a unique combination of barcodes (Ai, Bj) was introduced at the intersection of the microchannels (80 or 200 per chip), defining the spatial coordinate of each tissue spot. Nuclear integrity was then evaluated by DAPI staining, ensuring that nuclei remain morphologically intact throughout processing. Finally, barcoded Hi-C DNA fragments and biotinylated cDNA were released via reverse crosslinking. The cDNA was isolated using streptavidin beads and processed into an RNA-seq library via template switching, while the Hi-C DNA underwent splint ligation for DNA library construction. Sequencing both libraries yielded spatially resolved genomic and transcriptomic datasets from the same tissue section.

**Figure 1.**
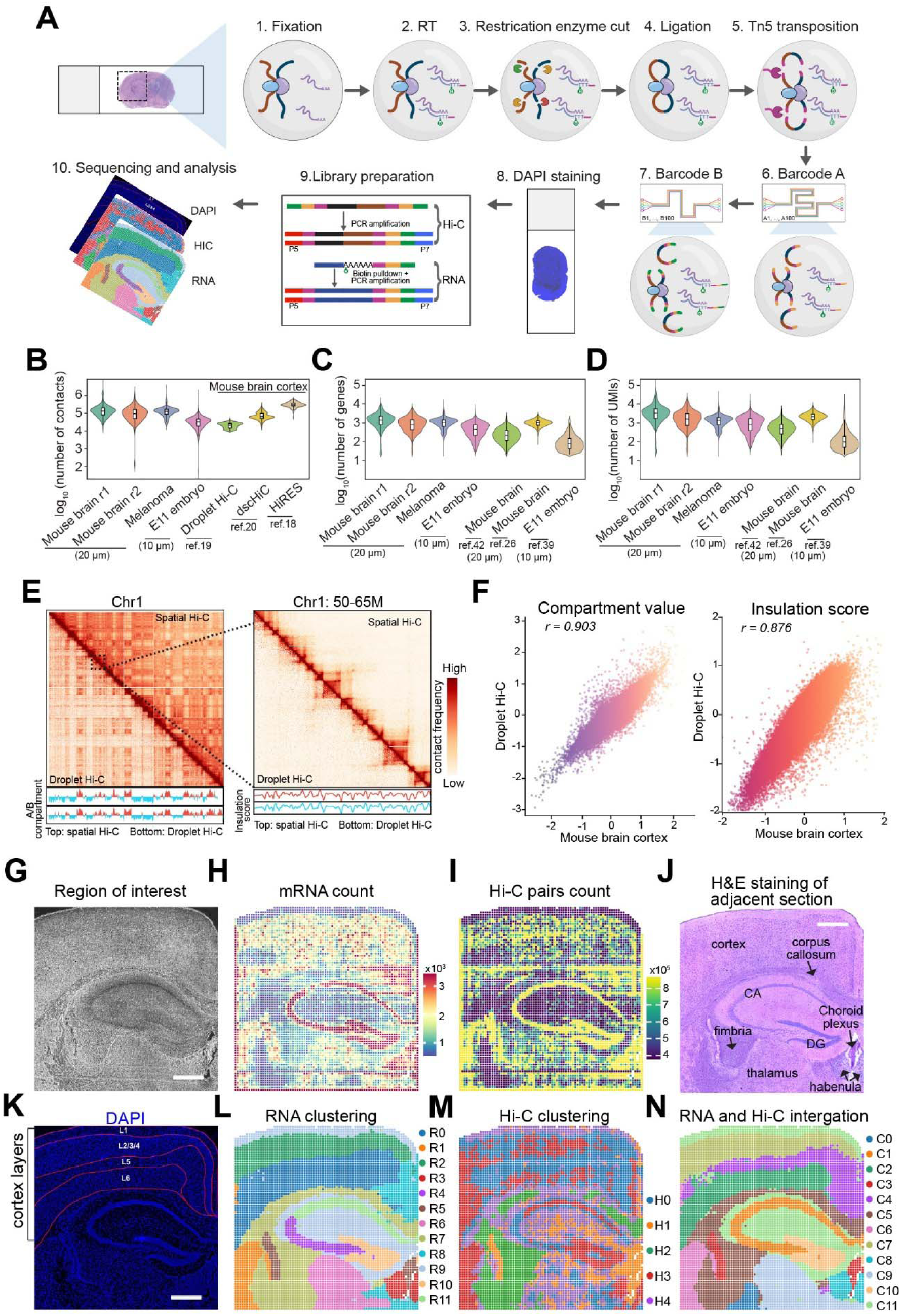
Overview, data quality assessment, and spatial mapping of the Spatial Hi-C-RNA method in mouse brain. (A) Workflow of Spatial Hi-C-RNA. Fresh-frozen tissue sections are subjected to (1) fixation, (2) reverse transcription, (3) restriction enzyme digestion, (4) proximity ligation, and (5) Tn5 transposition. Sequential barcoding is performed through (6) application of Barcode A, (7) Barcode B, and (8) DAPI staining, followed by (9) library preparation to generate spatially indexed RNA and Hi-C fragments. (10) Sequencing and downstream computational analysis yield co-registered maps of chromatin conformation, transcriptomes, and tissue architecture. (B-D) Quantitative metrics of Spatial Hi-C-RNA data quality in mouse brain compared with other benchmark datasets. Distribution of unique Hi-C contacts (B), RNA (C) and UMI (D) counts per pixel in different tissues at varying spatial pixel sizes. Violin plots illustrate the range, distribution, and median values for each dataset. (E) Chromosome-wide and zoomed-in contact matrices for Chr1 comparing Spatial Hi-C and droplet Hi-C. Left: full-chromosome 100-kb resolution maps. Right: zoomed 50–65 Mb region at 25-kb resolution. Bottom tracks are for A/B compartment and insulation score tracks. (F) Genome-wide comparison of compartment values (left) and insulation scores (right) between Spatial Hi-C and droplet Hi-C, showing strong correlation of large-scale and domain-level chromatin features across platforms. (G-K) Spatial distribution of molecular signal intensities across the tissue section. (G) Region of interest. Spatial RNA (H) and Hi-C (I) count maps. (J) H&E staining of an adjacent section. (K) API staining of the region of interest, highlighting cortical layer organization (L1–L6). Scale bar: 500 µm. (L-N) Spatial clustering and integrative analysis. (L) RNA-based clustering delineates major transcriptomic domains. (M) Hi-C–based clustering reveals chromatin-state-defined spatial compartments. (N) Joint RNA and Hi-C integration uncovers combined regulatory–structural tissue architectures, resolving coordinated transcriptomic and chromatin-state domains across the cortex.

We applied spatial Hi-C-RNA to diverse biological systems, including the adult mouse brains (2-months-old), human melanoma, and the embryonic day 11.5 (E11.5) mouse embryo, to benchmark performance across varying tissue types and resolutions. The mouse brain and melanoma were profiled using an 80 × 80 barcoding array (20 μm pixels, 3.2 × 3.2 mm area), while the E11.5 embryo was profiled with a high-density 200 × 200 array (10 μm pixels, 4 × 4 mm area) to resolve fine-scale spatial features.

We first assessed data quality in adult mouse brains. Spatial Hi-C-RNA generated 6,275 spatial pixels with a median of 593,680 unique read pairs per pixel, including 136,166 *cis* (>1 kb) and 42,457 *trans* contacts (41.9% duplicate rate; Table S1). The number of detected chromatin contacts was comparable to established Tn5-based single-cell Hi-C methods^18–20^ (Figure 1B). Simultaneously, the RNA modality detected a median of 1,458 genes and 3,210 UMIs per pixel, comparable to previous spatial transcriptomic studies^26,42^ (Figure 1C and 1D). An independent replicate (mouse brain r2) demonstrated high reproducibility in chromatin contact profiles and transcriptomic coverage (Figure 1C and 1D; Table S2). Furthermore, aggregate gene expression profiles showed strong concordance across biological replicates and with published datasets, confirming the fidelity of the RNA data^19,20^ (Figure S1A and S1B).

The captured chromatin architecture was similarly robust. Contact probability decay curves were highly consistent across samples (Figure S1C). To further benchmark performance, we compared Spatial Hi-C-RNA with single-cell droplet Hi-C.^19^ Our spatial approach captured a higher proportion of chromatin contacts (>1 kb) while maintaining comparable *trans* contact levels (Figure S1D). Aggregated pseudobulk contact maps, compartment scores, and insulation scores faithfully recapitulated known genome organization (Figure 1E). Correlation analyses revealed strong concordance between spatial Hi-C replicates and between spatial Hi-C and reference droplet Hi-C datasets (Figure 1F and S1E–L), underscoring the robustness and reproducibility of the platform for mapping 3D genome structure *in situ*.

### Spatial Hi-C–RNA links chromatin architecture to cell-type-specific gene regulation in the mouse brain

To investigate how transcriptional and chromatin architectures jointly shape spatial organization in the mouse brain, we generated spatial maps from the adult brain RNA and Hi-C data. Tissue morphology was visualized using DAPI staining on the profiled section and H&E staining on an adjacent section (Figure 1G, 1J, and 1K). Both mRNA expression patterns and Hi-C contact maps closely mirrored the underlying tissue architecture (Figure 1H, 1I, and S2A). Unsupervised spatial clustering (see Methods) of RNA modality identified twelve distinct transcriptional domains (R0–R11) that mapped precisely to anatomical regions (Figure 1L and S2B). In contrast, spatial clustering of chromatin features derived from Fast-Higashi^43^ embedding with CellCharter^44^ defined five Hi-C domains (H0–H4) with less distinct spatial boundaries (Figure 1M and S2B), consistent with previous observations that single-cell Hi-C resolves fewer cell types than single-cell RNA-seq.^13,21^ However, integrating both modalities revealed additional subregions defined by coordinated transcriptional and chromatin structural features (Figure 1N). For example, the cortical region was subdivided into multiple clusters, and distinct domains emerged within the thalamus. These results demonstrate that the synergy of RNA and Hi-C data enhances spatial resolution and uncovers combinatorial regulatory patterns that are not apparent from either modality alone.

We next asked whether known chromatin structural differences between neuronal and non-neuronal cells are preserved in a spatial context. Previous studies indicate that neurons exhibit a higher frequency of short-range chromatin interactions than non-neuronal cells.^13,22^ To quantify this spatially, we computed contact distance profiles from spatial Hi-C data, separating short- and long-range interactions (Figure 2A–C and S2C– D). Neuron-enriched regions—including cortex (except layer 1), hippocampal CA fields, and the dentate gyrus^13^—displayed higher short-to-long contact ratios compared with white matter–enriched regions (corpus callosum and fimbria),^45^ or the epithelial- and vascular-rich choroid plexus.^46^ Beyond differences in contact distance, global chromatin interaction profiles were highly concordant within neuron clusters and, separately, within non-neuronal clusters (Figure 2D). This structural divergence extended to higher-order folding, as evidenced by dynamic remodeling of A/B compartments between neuron- and non-neuron–enriched domains (Figure 2E and S2E). These spatial patterns likely reflect both intrinsic differences in chromatin organization and regional variation in the proportions of neuronal and non-neuronal cell types.

**Figure 2.**
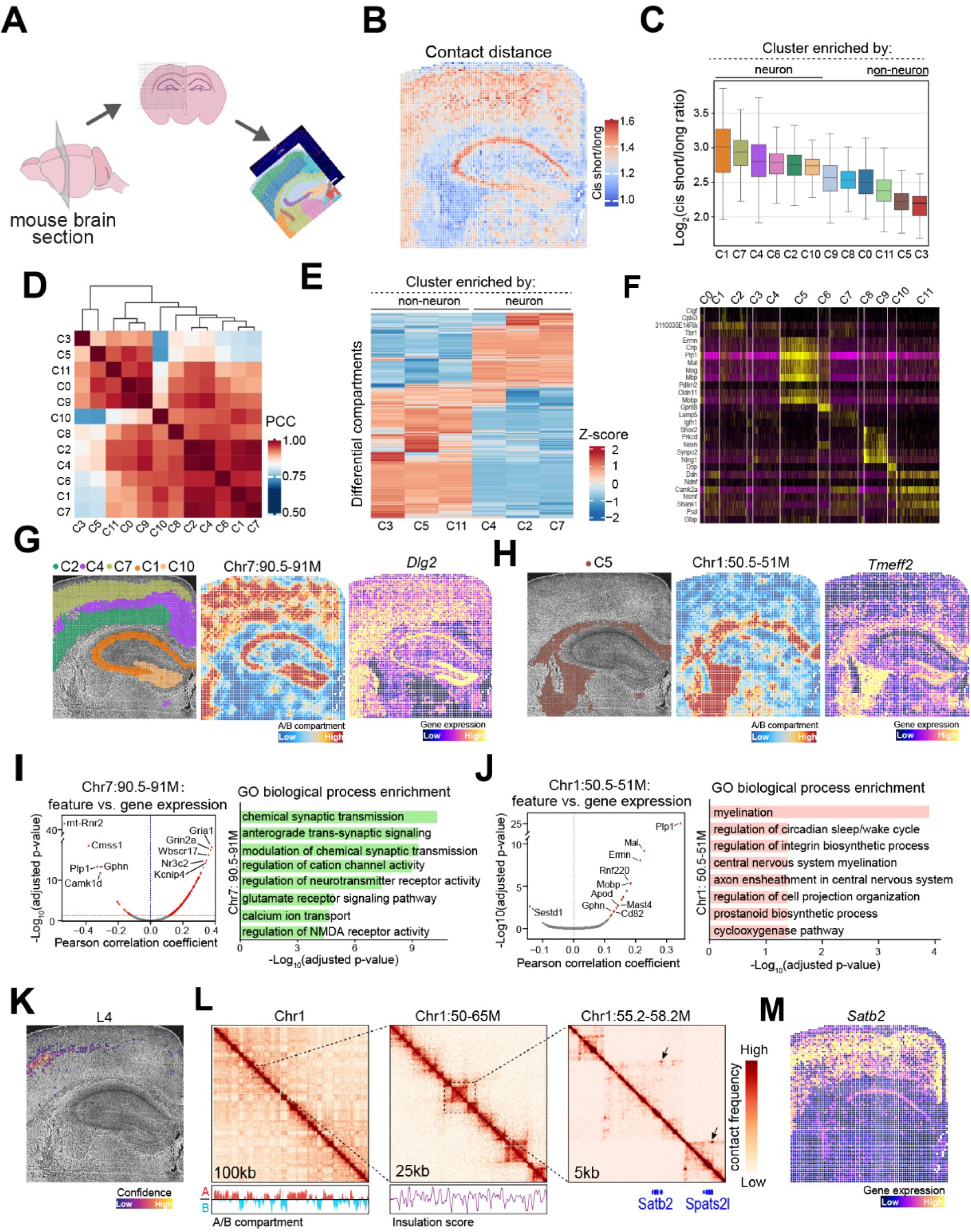
Spatial genomic compartmentalization and integration with gene expression in the mouse brain. (A) Schematic of Spatial Hi-C–RNA data processing in mouse brain cortex, illustrating extraction of tissue sections, spatial barcoding, and generation of co-registered chromatin and transcriptomic maps. (B-C) Spatial pattern of cis short/long contact distance across the mouse brain (B). Cluster-wise analysis of cis short/long ratio derived from Hi-C/RNA integrated clusters (C). (D) Pearson correlation coefficient (PCC) matrix showing similarity of A/B compartment profiles across spatially defined Hi-C/RNA integrated clusters. (E) Heatmap showing A/B compartment switches among selected Hi-C clusters, with scaled and sorted compartment scores. (F) Heatmap showing the normalized gene expression of markers corresponding to each Hi-C/RNA integrated cluster. (G-H) Spatial patterns of compartment scores (middle) and gene expressions (right) for representative domains (left): clusters C2, C4, C7, C1, and C10 represent neuron- enriched regions (G) and cluster C5 represents oligodendrocyte-enriched regions. (I-J) PCC analysis showing correlations between selected chromatin features and associated genes across spatial pixels, with go ontology enrichment summarizing biological processes linked to positively correlated genes. (K-M) Spatial pattern of L4 excitatory neurons (K). Heatmaps of L4 excitatory neurons at 100, 25, 5 kb resolutions. Bottom tracks are A/B compartment and insulation score tracks. *Satb1* locus associated a L4-specific loop (L), and its spatial gene expression (M).

Although chromatin A/B compartments are known to correlate with gene expression,^20–22^ how chromatin architecture regulates transcription in a spatially resolved manner remains underexplored. We first validated our spatial RNA-seq data using established markers, confirming the accurate cell identity assignments (Figure 2F and S2F). For example, *Mef2c* was highly expressed in the cortex, *Plp1* marked the corpus callosum, and *Camk2a* was enriched in cortical layer 1 and hippocampal regions (Figure S2G). Next, to identify the chromatin features underlying cluster-specific transcriptional programs, we applied a Random Forest classifier to A/B compartment scores derived from the spatial Hi-C data (see Methods), using spatial cluster identity as the prediction target. Feature importance values from the classifier were then used to prioritize chromatin bins that most effectively distinguished the spatial clusters. We focused on two top-ranking bins: Chr7:90.5–91 Mb, associated with neuron-enriched domains (clusters C1, C2, C4, C7, C10), and Chr1:50.5–51 Mb, associated with oligodendrocyte- dominated domain (cluster C5). As expected, the compartment states of these bins showed strong spatial concordance with the anatomical layout of their respective domains, and the expression of corresponding marker genes (*Dlg2* and *Tmeff2*) mirrored these spatial distributions (Figure 2G and 2H). Building on the principle that patterns of gene–gene^47^ and peak–peak^48^ co-expression often capture higher-order pathway activity, we hypothesized that A/B compartment states might similarly covary with distal gene expression, suggesting coordinated spatial regulation beyond local associations. We therefore further analyzed these selected bins to evaluate potential long-range correlations. The neuronal bin at Chr7:90.5–91 Mb (Figure 2I), selectively activated in cortical L2–6 and hippocampal CA/DG neurons, correlated strongly with the expression of *Gria1*, *Grin2a*, *Wbscr17*, *Nr3c2*, and *Kcnip4*. Gene Ontology (GO) analysis revealed enrichment for excitatory neurotransmission pathways (see Method), including chemical synaptic transmission, anterograde trans-synaptic signaling, glutamate receptor signaling, and calcium/cation transport, as well as regulation of NMDA receptor activity. Conversely, the oligodendrocyte-associated bin at Chr1:50.5– 51 Mb (Figure 2J) correlated with canonical myelination genes, including *Plp1*, *Mal*, *Ermn*, *Rnf220*, and *Apod*. These genes were enriched for biological processes essential to oligodendrocyte function, such as myelination, central nervous system axon ensheathment, and regulation of cell projection organization. These findings indicate that spatially resolved A-compartment activation serves as a reliable proxy for coordinated transcriptional modules and their associated biological functions.

To assess the resolution of spatial Hi-C-RNA at the scale of chromatin loop (5 kb), we first deconvolved cell types within each spatial cluster using TACCO,^49^ an optimal transport–based framework that integrates our data with external single-cell RNA-seq references^45^ from the adult mouse brain (see Methods). This approach resolved major neuronal and non-neuronal populations (Figure S3A) and revealed a distinct spatial distribution of astrocytes, excitatory neurons (L2/3 IT, L4, L6 CT), inhibitory neurons (Pvalb, Sst), and oligodendrocytes (Figure S3B and S3C). Cell type assignments were further validated by canonical marker genes, including *Slc1a3* (astrocytes), *Cux2* (L2/3 IT neurons), *Rorb* (L4 neurons), *Tle4* (L6 CT neurons), *Mbp* (oligodendrocytes), *Pvalb* (Pvalb interneurons), and *Zic1* (Sst neurons) (Figure S3B and S3C). By aggregating Hi-C reads from spatially defined cell populations, we recovered clear chromatin loops at key marker loci. For example, we observed strong chromatin loops at the *Satb2* locus specifically in L2/3 IT and L4 excitatory neurons (Figure 2L and S4A), concordant with the high expression of *Satb2* in the same cell types (Figure 2M). Similarly, the *Spats2l* locus displayed prominent looping interactions and elevated gene expression in the same excitatory neuron populations (Figure 2L, S4A, and S4B). In contrast, these loops were absent in astrocytes and oligodendrocytes, where gene expression was minimal (Figure S4A and S4B). Taken together, these results validate that Spatial Hi-C-RNA reliably captures multiscale chromatin organization—from compartments to loops—and directly links these features to cell-type-specific transcriptional programs within intact tissues.

### 3D genome rewiring and transcriptional reprogramming in the tumor microenvironment

To investigate how 3D genome architecture shapes the complex heterogeneity of the tumor microenvironment (TME), we applied Spatial Hi-C–RNA to human melanoma (Figure 3A). We first guided our region of interest (ROI) selection using S2Omics,^50^ an automated histology-based segmentation tool, which partitioned the H&E-stained tissue into ten distinct clusters representing diverse histological architectures (Figure 3B and 3C). We selected a region exhibiting high histological diversity (Figure 3B, red box) for subsequent spatial profiling and established a comprehensive multimodal spatial analysis pipeline. By integrating Spatial Hi-C-RNA (20 μm resolution; 3.2 × 3.2 mm) with spatial ATAC-seq profiled on an adjacent section (20 μm resolution; 4 × 4 mm), we constructed a spatially registered multi-omics map capturing chromatin conformation, chromatin accessibility, and transcriptional activity within the intact TME (Figure 3D, S5A, and S5B).

**Figure 3.**
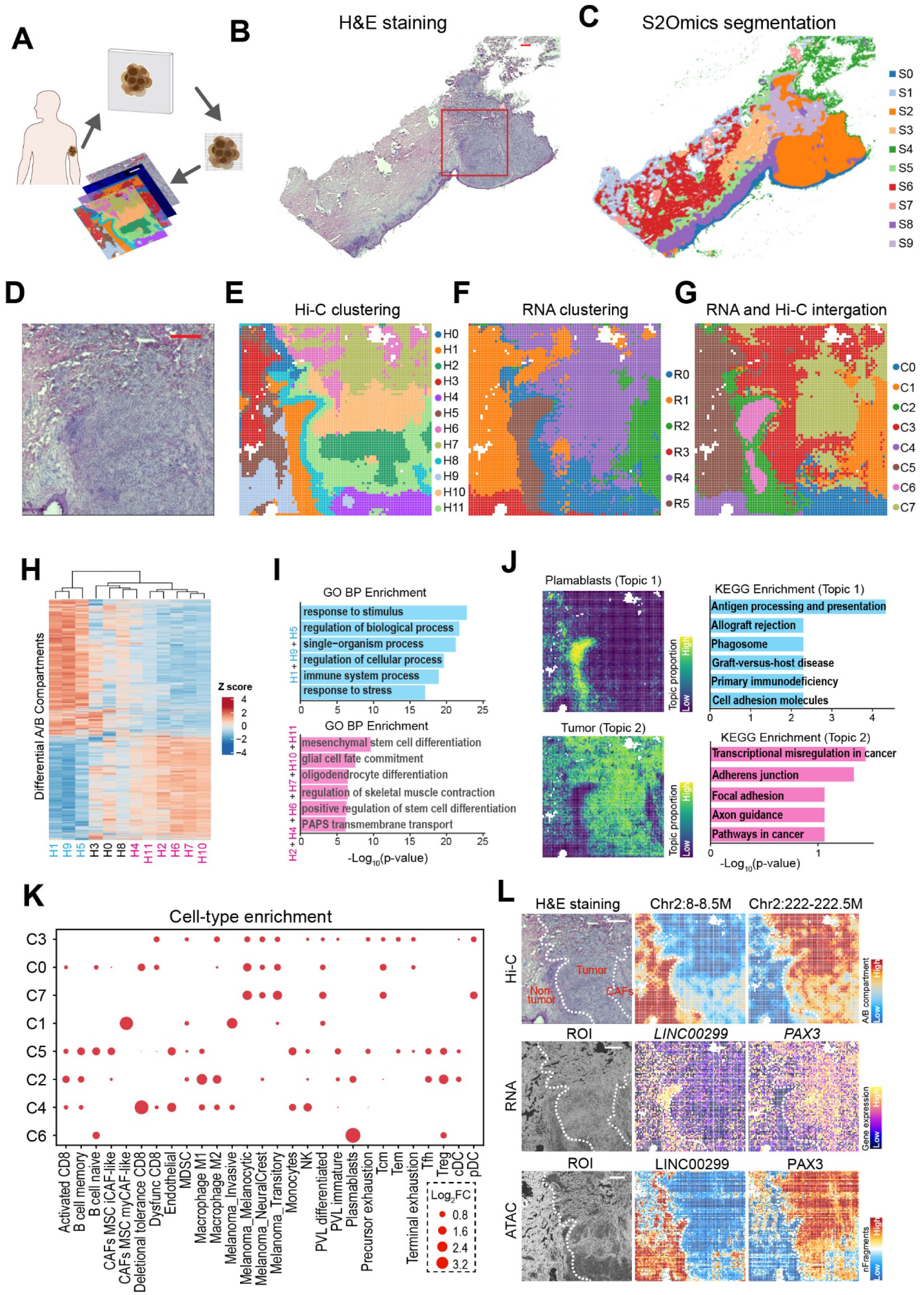
Spatial Hi-C–RNA integration reveals multicellular organization and chromatin architecture in human melanoma tissue. (A) Schematic of the Spatial Hi-C–RNA workflow on intact tumor sections. (B) H&E staining of the representative melanoma tissue section. Scale bar: 500 μm. (C) Automated S2Omics histology-based segmentation delineates ten distinct tissue architectures. (D) Zoom-in of the region of interest highlighting heterogeneous tumor microenvironments. Scale bar: 500 μm. (E) Spatial Hi-C clustering revealing chromatin architecture–based domains across the section. (F) Spatial RNA clustering identifies transcriptionally distinct regions. (G) Joint integration of Hi-C and RNA data defines multimodal clusters. (H) Heatmap showing A/B compartment switches among Hi-C clusters, with scaled and sorted compartment scores. (I) Gene Ontology (GO) highlights functional specialization for representative groups: top includes clusters H1, H5, and H9; bottom includes clusters H1, h2, H4, H6, H7, H10, and H11. (J) Spatial distributions of topics identified with spatial RNA-seq data (left), corresponding to plasmablasts (Topic 1) and tumor (Topic 2). KEGG pathway enrichment analysis for each topic (right). (K) Cell-type enrichment across each Hi-C/RNA integrated cluster (*n* = 8 clusters). (L) Multimodal visualization of selected loci (*LINC00299* and *PAX3*) showing concordant variation in chromatin contacts (Hi-C, top), gene expression (RNA, middle), and chromatin accessibility (ATAC, bottom) within anatomically defined regions (left). Scale bar: 500 μm.

We validated the performance of this multimodal approach in the complex tumor tissue. The Hi-C modality yielded a median of 652,994 unique read pairs per pixel (6,199 spatial pixels), capturing a rich landscape of chromatin interactions (median 132,974 *cis* (>1 kb) and 57,756 *trans* contacts per pixel; 43.2% duplicate rate; Table S3). Simultaneously, the RNA modality detected a median of 1,107 genes and 1,389 UMIs per pixel. In parallel, spatial ATAC-seq profiling from an adjacent tissue section produced a median of 39,991 unique fragments per pixel (Figure S5C). Unsupervised spatial clustering of the ATAC-seq data identified 5 spatially organized clusters (see Methods) (Figure S5D), providing complementary insights into chromatin accessibility patterns within the TME.

To investigate the spatial relationship between chromatin organization and transcriptional programs in the melanoma sample, we first performed independent spatial clustering of Hi-C and RNA modalities, followed by joint integration analysis. While RNA profiles identified 6 broad transcriptional clusters (R0–R5), Hi-C-based spatial clustering revealed 12 distinct spatial domains (H0–H11), capturing fine-scale variation in local chromatin conformation (Figure 3E and 3F). The finer partitioning in Hi-C data indicates that chromatin architecture captured layers of spatial heterogeneity— potentially reflecting epigenetic plasticity—that are not fully resolved by the transcriptome alone. Integrating both modalities yielded eight joint spatial clusters (C0– C7), which combined the structural sensitivity with the functional specificity of transcriptomic profiles (Figure 3G). To understand the genome organization underlying spatial heterogeneity, we analyzed A/B compartment switching across the Hi-C clusters (Figure 3H). Hierarchical clustering identified differentially compartmentalized regions and GO analysis of genomic regions with switched compartment states revealed two distinct functional groups. The first group (H1, H3, and H5) exhibited active compartment profiles enriched for pathways related to response to stimulus, stress, and immune activation (Figure 3I, top). In contrast, the second group (H2, H4, H6, H7, H10, and H11) was characterized by developmental and lineage-specifying programs, including mesenchymal differentiation, glial cell fate commitment, and stem cell regulation (Figure 3I, bottom). These patterns suggest that tumor chromatin remodeling involves both activation of stress-responsive networks and reconfiguration of developmental trajectories within the TME. Furthermore, the second group exhibited a higher short-to-long contact ratio (Figure S5E and S5F), consistent with a more compact chromatin configuration often observed in malignant states.^51^

We leveraged our spatial RNA modality to map the cellular context of these structural domains. Using Starfysh^52^ for reference-free deconvolution (see Methods), we resolved the spatial distribution of melanoma cells, stromal components, and immune populations (Figure S6A). We identified distinct melanoma niches—melanocytic, neural crest–like, and transitory states—bordered by cancer-associated fibroblasts (CAFs) and plasmablasts (Figure S6B). Expression of canonical markers confirmed these annotations, with *IGHG1* marking plasmablasts,^53^ *OCA2* labeling melanocytic melanoma cells,^54^ and *COL1A1* delineating fibroblast-rich areas^55^ (Figure S6C). To characterize spatial transcriptional programs, we applied STAMP^56^ to the spatial RNA modality (see Methods), which revealed multiple spatially coherent gene expression topics across the melanoma section. Representative topics corresponding to plasmablasts, tumor cells, keratinocytes, and CAFs are shown in Figure 3J, S6D, and S6E. The plasmablast-associated topic was enriched for immune pathways, including antigen processing and presentation, whereas the tumor topic highlighted pathways such as transcriptional regulation in cancer and focal adhesion (Figure 3J). Keratinocyte-associated regions were enriched for immune and stress-response pathways (Figure S6D). In contrast, the CAF-associated topic displayed strong enrichment for extracellular matrix remodeling and cell-matrix interaction pathways, including ECM–receptor interaction and focal adhesion (Figure S6E). Cell-type enrichment analysis further revealed distinct cellular compositions across the integrated Hi-C/RNA clusters (Figure 3K), complementing the spatial transcriptional topics identified above. Clusters C0, C3, and C7 were enriched for melanoma cells, whereas C2 and C6 were dominated by immune populations, and C1 displayed strong CAF signatures. Spatial mapping of these clusters showed a coherent tissue architecture with clear boundaries demarcating the tumor, CAF, and surrounding non-tumor regions (Figure 3L).

To examine how chromatin structure relates to transcriptional regulation across these spatial domains, we evaluated loci such as *LINC00299* and *PAX3*, the latter of which is a well-established melanoma-associated transcription factor.^57^ These loci exhibited contrasting regulatory landscapes across tumor and non-tumor regions. Hi-C data revealed that these loci occupied opposite A/B compartments, reflecting region-specific chromatin folding patterns (Figure 3L, top). Consistent with this, spatial RNA profiling showed elevated *PAX3* expression in tumor and CAF regions, whereas *LINC00299* was repressed in the same domains (Figure 3L, middle). Spatial ATAC-seq further supported these observations: *PAX3* displayed locally increased chromatin accessibility in tumor regions, whereas *LINC00299* remained comparatively inaccessible (Figure 3L, bottom). We next examined whether fine-scale 3D chromatin architecture could further distinguish malignant from non-malignant subregions. High-resolution Hi-C maps revealed a prominent chromatin loop at the *PAX3* locus that was specific to tumor and CAF clusters (C7 and C1) but largely absent in non-tumor regions (C4 and C6) (Figure S7A). A similar pattern was observed at the *ACSL3* locus, which showed tumor-specific chromatin loops, active compartment status, elevated chromatin accessibility, and increased expression in tumor and CAF clusters (Figure S7A and S7B). Together, these results demonstrate a coordinated relationship between multiscale 3D genome organization, chromatin accessibility, and transcriptional programs within the spatially organized melanoma microenvironment.

### Integrative multimodal profiling reveals chromatin–gene coupling underlying melanoma progression

To link chromatin structural variation with tissue-level organization, we examined how spatial cellular architecture and chromatin dynamics differ between the CAF-enriched (C1) and tumor-dominant (C7) regions. Differential neighborhood enrichment (NE) analysis^44^ revealed distinct microenvironmental contexts between these clusters (Figure 4A). The CAF region exhibited strong spatial associations with immune populations, including activated and exhausted CD8⁺ T cells, B cells, and macrophages, consistent with an immune-infiltrated stromal interface. In contrast, the tumor region was preferentially associated with melanoma cell populations, particularly invasive and melanocytic subtypes, indicating a tumor-dominant niche with limited immune infiltration. To reconstruct the molecular trajectories driving these spatial states, we performed pseudotime analysis based on both the Hi-C and RNA profiles (Figure 4B and 4C). Both modalities resolved a continuous transition from melanocytic to invasive melanoma states, reflecting coordinated chromatin remodeling and transcriptional reprogramming along tumor progression.

**Figure 4.**
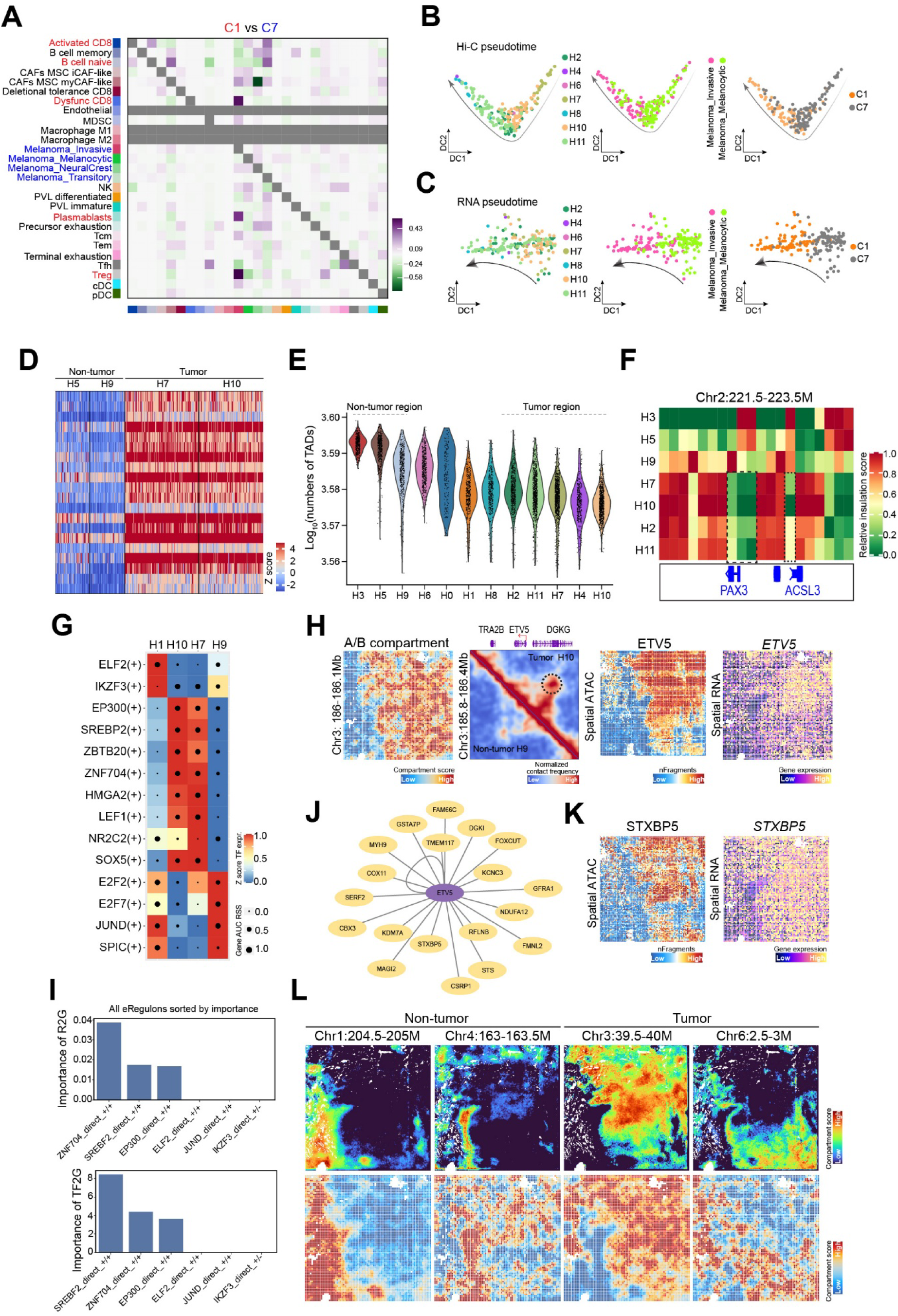
Integration of Spatial Hi-C–RNA and ATAC-seq reveals chromatin rewiring, transcriptional programs, and regulatory network remodeling in human melanoma. (A) Cluster NE (neighborhood enrichment) analysis between two spatial clusters C1 and C7 in human melanoma. Examples of differentially enriched neighborhoods are colored. (B-C) Inferred melanoma_invasive lineage trajectory from Spatial Hi-C (B) and RNA (C) profiles, displayed by Hi-C clusters (left), annotated cell types (melanoma_melanocytic and melanoma_invasive; middle), and Hi-C/RNA integrated clusters (C1 and C7; right). (D) Heatmap showing subcompartment assignments (Z-score normalized) for selected non-tumor (H5, H9) and tumor (H7, H10) Hi-C clusters. (E) Violin plots showing the distribution of topologically associating domain (TAD) counts (log_10_-transformed) across individual Hi-C clusters. (F) Aggregated cluster-specific insulation score (100 kb resolution) at the *PAX3* and *ACSL3* locus. (G) Heatmap/dot-plot showing transcription factor expression of the eRegulon on a color scale and selected cluster specificity of the eRegulon on a size scale. (H) Spatial A/B compartment map demonstrating disruption of normal compartment organization in melanoma regions (left). Chromatin loop of *ETV5* between tumor cluster H10 and non-tumor cluster H9 at 5 kb resolution. Right: spatial chromatin accessibility and gene expression maps of *ETV5*. (I) Region-to-gene (top) and transcription factor-to-gene (top) interactions of ETV5 across the whole dataset. (J) Gene regulatory network centered on transcription factor ETV5 inferred from spatial ATAC/RNA data. (K) Spatial chromatin accessibility and gene expression of ETV5-GRN target STXBP5. (L) iStar super-resolution predictions of spatial A/B compartments closely recapitulate original spatial Hi-C measurements across representative loci in non-tumor and tumor regions, preserving global domain structure while revealing finer chromatin variation aligned with local histopathology. Top: iStar-predicted super-resolution maps; bottom: original-resolution compartment scores.

Beyond global A/B compartmentalization, the 3D genome can be further resolved into finer subcompartments that more precisely capture chromatin activity states.^58^ To explore how these fine-scale structures are altered in melanoma, we analyzed a 4-Mb region on chromosome 5 (Chr5:123.5–127.5Mb) that consistently resides within active A compartments in tumor regions (Figure S7C). Application of the scGHOST^59^ framework revealed a marked redistribution from repressive subcompartments (scB1–scB3) toward active subcompartments (scA1 and scA2) in tumor Hi-C clusters (Figures 4D and S7D) (see Method). At a higher resolution (100 kb), the tumor region exhibited a reduced number of topologically associating domains (TADs) compared to non-tumor regions (Figure 4E), indicative of boundary erosion and increased inter-domain connectivity (see Methods). This global relaxation of chromatin architecture was exemplified at the *PAX3*– *ACSL3* locus, which displayed diminished insulation scores and enhanced local interactions in tumor regions (Figures 4F, S7A, and S7B), suggesting that activation of oncogenic programs is facilitated by a loss of structural constraints.

We next sought to identify enhancer-driven regulons (eRegulons) driving chromatin architectural changes. Joint SCENIC+^60^ analysis of spatial ATAC-seq and RNA-seq data revealed region-specific regulators of infiltrating immune populations, such as ELF2 and IKZF3, as well as tumor-region regulators like LEF1 and HMGA2, with spatial profiles further validating their activity (Figure 4G and S8A). To further investigate how these eRegulons regulate the downstream targets, we focused on ETV5, a transcription factor overexpressed in melanoma compared to normal skin.^61^ Examination of the *ETV5* locus revealed tumor-specific activation, with strong A-compartment activity, prominent chromatin looping, increased chromatin accessibility, and elevated gene expression (Figure 4H). We used ETV5 as an example to examine the influence of eRegulons on chromatin accessibility and gene expression. In Figure 4I, we assessed region-to-gene and transcription factor-to-gene interactions for ETV5 across the entire dataset. Our analysis identified ZNF704, SREBF2, and EP300 as the most prominent tumor-specific eRegulons associated with ETV5, while non-tumor-specific eRegulons, including ELF2, JUND, and IKZF3, showed minimal impact (Figure 4I). These findings aligned with Figure 4G, where ZNF704, SREBF2, and EP300 were also identified as key regulators in tumor regions. To illustrate the downstream consequences of ETV5 activation, we highlighted *STXBP5* as a representative target. *STXBP5* locus exhibited tumor-enriched chromatin accessibility and RNA expression, demonstrating coordinated activation of the ETV5-centered regulatory program (Figure 4J and 4K).

Finally, we assessed the potential of spatial Hi-C for resolving fine-scale, exactly single-cell resolution and continuous image-like chromatin-architecture activity. We applied iStar^62^, a deep learning framework that integrates high-resolution histological features (H&E) to predict super-resolved molecular maps (see Methods). Comparison of iStar- predicted A/B compartment scores with measured spatial Hi-C compartment maps across representative loci showed that the iStar-predicted maps preserved global domain organization while resolving fine-scale local variations closely aligned with tumor histopathology (Figure 4L). These results highlight the feasibility of combining spatial Hi-C with image-derived features to achieve high-accuracy inference of fine-scale chromatin organization. This approach also holds promise for clinical applications, potentially enabling the precise estimation of higher-resolution 3D genome architecture directly from standard, low-cost H&E sections for comprehensive molecular pathology and precision diagnostics.

In summary, the application of Spatial Hi-C-RNA to human melanoma revealed a spatially organized interplay between chromatin architecture and transcriptional regulation within the tumor microenvironment. By integrating Hi-C, RNA, and ATAC modalities, we uncovered the coordinated chromatin remodeling and transcriptional reprogramming associated with melanoma progression. Together, these findings established Spatial HiC-RNA as a powerful framework for decoding the physical and functional logic of gene regulation in complex tissues.

### Spatial 3D genome and transcriptome mapping defines chromatin state dynamics during embryonic development

To jointly visualize genome organization and transcriptional activity during embryogenesis, we applied Spatial Hi-C-RNA to an E11.5 mouse embryo section at near single-cell resolution (10 µm) (Figure 5A). DAPI staining confirmed tissue integrity (Figure S9A). In total, 32,799 spatial pixels were captured, yielding a median of 125,582 unique Hi-C read pairs per pixel, comprising 34,816 *cis* (>1 kb) and 26,508 *trans* contacts (Table S4). Simultaneously, the RNA modality detected a median of 453 genes and 869 UMIs per pixel (Figure S9B and S9C). Unsupervised clustering of each modality revealed distinct spatial organizations of transcriptional and chromatin states (Figure 5B and 5C). RNA profiles resolved 14 transcriptionally defined clusters corresponding to major embryonic anatomical territories (Figure 5B), whereas Hi-C data defined 6 broader spatial domains (Figure 5C). Integrating both datasets yielded 12 refined multimodal clusters (Figure 5D) that aligned precisely with anatomical structures. Representative marker genes, such as *Dpp10* (craniofacial epithelium), *Pax3* (neural progenitors), *Hbb-bt* (erythroid lineages), and *Adamts18*/*Col2a1* (mesenchyme/cartilage), validated the biological specificity of these domains (Figure S9D).

**Figure 5.**
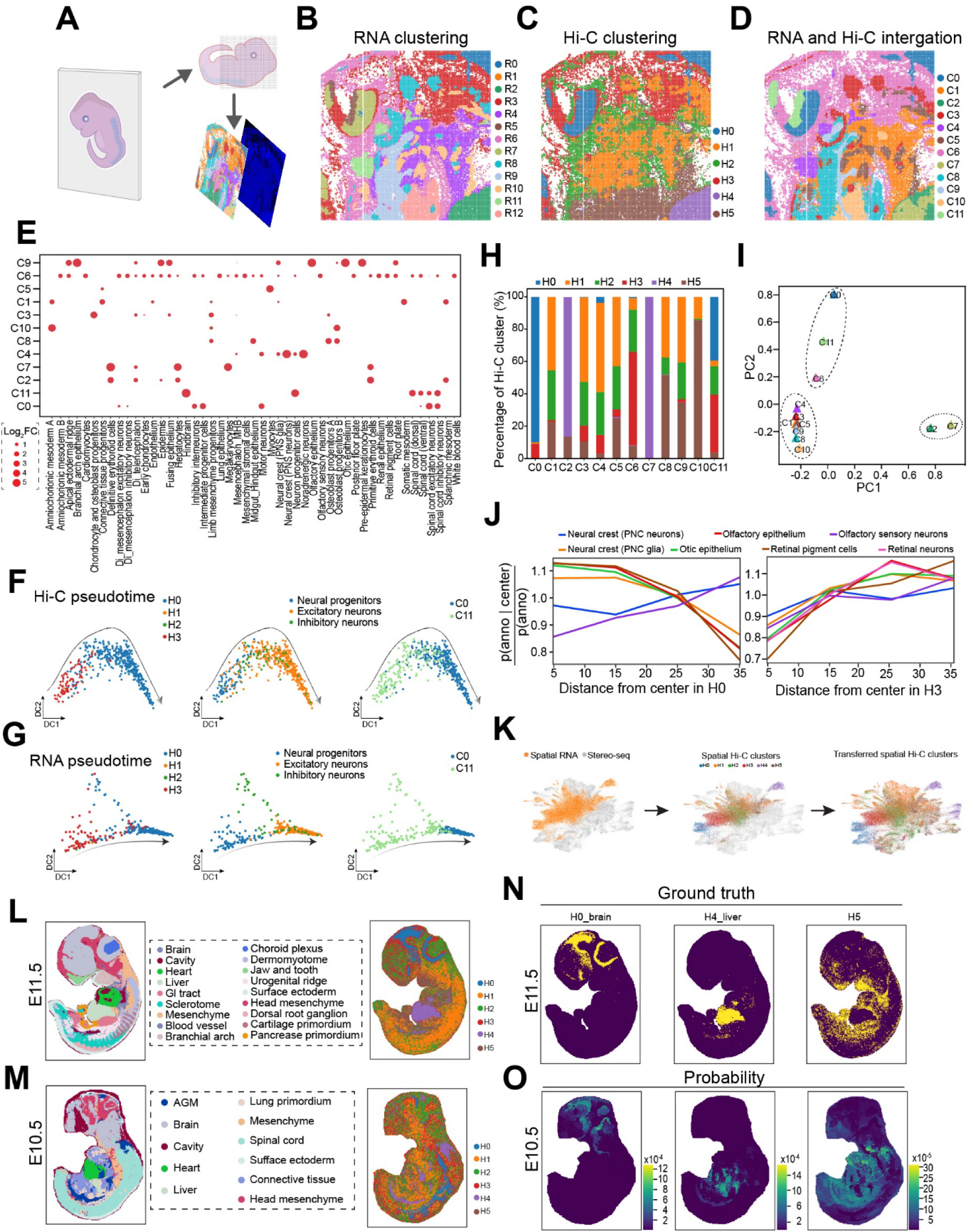
Spatial co-profiling of 3D genome architecture and gene expression in the mouse embryo. (A) Overview of spatial Hi-C–RNA profiling of an E11.5 mouse embryo section at 10-µm resolution. (B-D) Unsupervised clustering of spatial RNA expression (B) and chromatin contact maps (C), each producing spatially coherent domains corresponding to major anatomical regions. (D) Joint embedding of Hi-C and RNA profiles yields 12 refined multimodal clusters with improved alignment to embryonic structures. (E) Cell-type enrichment of integrated Hi-C/RNA clusters, derived by mapping spatial transcriptomes to a single-cell RNA-seq reference. (F-G) Pseudotime trajectories reconstructed independently from spatial Hi-C (I) and RNA (J) datasets, displayed by Hi-C clusters (left), annotated cell types (neural progenitors and excitatory/inhibitory neurons; middle), and Hi-C/RNA integrated clusters (C0 and C11; right). (H) Distribution of Hi-C structural domains across Hi-C/RNA-integrated clusters (*n* = 12 clusters), highlighting mixed chromatin states across the tissue section. (I) PCA showing distinct segregation of Hi-C clusters. (J) Co-occurrence analysis quantifying the spatial arrangement of cell types surrounding neural progenitors from the center in cluster H0 (left) and H3 (right), revealing proximal enrichment of early neurodevelopmental lineages and peripheral localization of sensory and epithelial derivatives. (K) Workflow for transferring Hi-C cluster labels onto Stereo-seq datasets using scVI-based integration and k-nearest neighbor classification. (L-M) Projected Hi-C clusters mapped onto E11.5 (L) and E10.5 (M) Stereo-seq transcriptomes (left), recapitulating major anatomical territories (right). (N-O) Cross-stage projection of Hi-C chromatin states onto E11.5 (N) and E10.5 (O) embryos, revealing organ-specific chromatin maturation, including early activation in liver (H4), relative stability in brain-associated domains (H0, H3), and moderate transitions in heart (H5).

To map cellular composition, we deconvolved spatial RNA profiles using a single-cell RNA-seq reference^63^ via TACCO (Figure 5E and S9E). Most spatial clusters exhibited selective enrichment for distinct cell types, reflecting emerging tissue-specific microenvironments (Figure 5E). For example, clusters C11 and C0 were enriched for neural progenitors and differentiating neurons; cluster C3 contained abundant chondrocyte and osteoblast progenitors, distinguishing developing skeletal elements; C2 and C7 clusters exhibited high representation of erythroid lineages; and C4 and C8 encompassed epithelial and surface ectoderm derivatives, including otic and olfactory lineages. These annotations confirm that multimodal clusters recapitulated major developmental compartments with accurate spatial localization. Focusing on neural progenitors and differentiating neurons, we reconstructed developmental trajectories using both Hi-C and RNA profiles (Figure 5F and 5G). Both modalities converged on a coherent pseudotemporal progression from progenitors to excitatory and inhibitory neurons, highlighting the coordinated reorganization of chromatin architecture and transcriptional changes during neuronal maturation. In parallel, we observed an increase in the *cis* short/long contact ratio in the ventral hindbrain (Figure S10A), indicating progressive changes in genome organization as cells differentiate.

While some Hi-C domains displayed clear regional specificity, others appeared less distinct (Figure 5H and 5I), suggesting that 3D genome architecture does not fully resolve cell-state complexity at this developmental stage^18^. Nevertheless, this structural organization captures latent developmental heterogeneity within morphologically similar regions. For instance, the neural progenitor-enriched cluster C11 was composed of a mixture of Hi-C clusters H0 and H3 (Figure 5H). This indicates that distinct chromatin states coexist within the same coarse-grained tissue region. Since our trajectory analysis identified H3 as an earlier developmental state and H0 as a later state (Figure 5F), these intra-tissue chromatin variations may delineate spatially distinct niches that correspond to different stages of developmental progression. We further examined cell-type co-occurrence patterns surrounding neural progenitor cells in the H0 and H3 Hi-C domains using TACCO-based co-occurrence analysis (Figure 5J).^49^ In the early-state H3 domain, neural progenitors showed high co-occurrence with neural crest cells—a lineage closely linked to early neurogenesis—regardless of distance. In contrast, in the late-state H0 domain, neural progenitors showed stronger local co-occurrence with specialized sensory and epithelial populations (retinal/olfactory/otic), consistent with the maturation of peripheral sensory structures. These findings highlight the ability of spatial Hi-C to resolve subtle niche-level heterogeneity, capturing distinct cell–cell interaction patterns that arise from developmental stage differences within the same tissue structure.

To characterize the structural basis of these developmental transitions, we analyzed chromatin architecture across multiple scales. Globally, we observed a progressive increase in the *cis* short-to-long contact ratio in the ventral hindbrain (Figure S10A), indicating that neuronal differentiation is accompanied by a global reorganization of the genome. Regionally, we analyzed spatially resolved A/B compartment patterns. Along chromosome 19, several genomic regions showed increasing A-compartment features in maturing neuronal populations (Figure S10B), coinciding with the upregulation of nearby neural-associated genes such as *Sorcs3* and *Sorcs1* (Figure S10C). At finer scales (5 kb), we assessed gene-specific chromatin interactions. Genes enriched in differentiating neurons, such as *Dscaml1*, exhibited strengthened loop contacts within neuronal domains (Figure S10D), whereas the progenitor marker *Racgap1* showed enhanced interactions in progenitor regions (Figure S10E). These examples illustrate that stage-specific gene regulation is accompanied by dynamic, spatially patterned remodeling of 3D genome architecture.

Finally, to further explore chromatin dynamics during organogenesis, we extended our analysis across spatiotemporal scales by integrating our E11.5 Spatial Hi-C-RNA data with high-resolution Stereo-seq transcriptomes from E10.5 and E11.5 embryos^64^ (see Methods). To achieve robust integration across modalities, we first aligned the datasets into a shared latent space using scVI^65^, which corrects for batch effects and modality-specific technical noise (Figure 5K). A k-nearest neighbor (kNN) classifier trained on Hi-C clusters was then used to assign corresponding cluster identities to the Stereo-seq spots (Figure 5K,5L, and 5M). The projected clusters displayed distinct, spatially coherent patterns across the embryo. For example, H0 was highly enriched in the developing brain and H4 in the liver, demonstrating that chromatin domain information can be effectively inferred from spatial transcriptomic data. We then applied Moscot^66^ to project the chromatin-defined cluster identities onto the spatiotemporal transcriptomic maps from Stereo-seq (E10.5 and E11.5 embryos; see Methods). This analysis revealed pronounced organ-specific differences in the pace of chromatin remodeling (Figure 5N, 5O, and S10F). In the developing liver, the proportion of the H4 chromatin signature remained stable between E10.5 and E11.5, suggesting that the global chromatin state of hepatic tissue is maintained during this window. In contrast, the developing brain exhibited shifts in chromatin state composition, with the proportions of clusters H0 and H3 changing markedly between stages, reflecting the rapid and extensive chromatin reorganization accompanying neurogenesis. Additionally, the heart region displayed an increase in the H5 chromatin signature, indicative of ongoing cardiac maturation. These results highlight asynchronous chromatin reorganization across organs, reflecting distinct developmental trajectories during early embryogenesis.

## DISCUSSION

Gene expression in complex tissues is governed by a hierarchical 3D genome organization, ranging from local chromatin accessibility and histone modifications to higher-order structures such as loops, TADs, and compartments. While recent advances in spatial multi-omics technologies have enabled the *in situ* profiling of epigenetic and regulatory layers—such as DNA methylation, chromatin accessibility, and histone modifications^25,26,42^—the spatial principles linking 3D genome folding to transcriptional programs within intact tissues have remained largely uncharacterized. Here, we address this gap by developing Spatial Hi-C-RNA, a high-resolution microfluidic *in situ* barcoding strategy that simultaneously maps chromatin architecture and transcriptomes with their native tissue context. Across the adult mouse brain, human melanoma, and developing mouse embryo, this technology generated high-quality multimodal maps that aligned with established single-cell and spatial references, providing an integrated view of how the 3D genome orchestrates cellular function *in situ*.

By co-profiling 3D genome architecture and gene expression *in situ*, Spatial Hi-C-RNA reveals how structural and transcriptional programs jointly specify cellular identity. In the adult mouse brain, determining cell type solely by RNA overlooks the underlying regulatory potential, and our integrated clustering resolved fine anatomical domains and identified cell-type-specific chromatin features, such as neuronal- and oligodendrocyte-enriched A-compartment activations coordinated with transcriptional modules. In addition, Spatial Hi-C-RNA demonstrates that 3D genome rewiring is a spatially coherent mechanism of tumor evolution. In melanoma, we observed a tumor-specific reorganization characterized by boundary erosion, subcompartments switching, and the formation of distinct loops at oncogenic gene loci such as *PAX3* and *ACSL3*, linking these structural alterations to transcriptomic and chromatin-accessibility states. Pseudotemporal analyses further demonstrated that chromatin remodeling parallels transcriptional transitions from melanocytic to invasive tumor states, underscoring a tight coupling between genome architecture and tumor progression. Furthermore, this approach enables the reconstruction of spatiotemporal dynamics in developing tissues. In the mouse embryo, Hi-C and RNA trajectories converged to reveal the coherent lineage maturation, marked by the progressive A-compartment activation and strengthening of neuronal loops. Transferring Hi-C-defined chromatin states onto high-resolution Stereo-seq maps revealed asynchronous nature of organogenesis, with hepatic chromatin states stabilized early, neural chromatin architectures underwent protracted remodeling.

Beyond biological discovery, Spatial Hi-C-RNA represents a flexible and scalable framework for the next generation of spatial biology. Its deterministic barcoding architecture supports tunable resolutions (10–20 μm) and capture areas (up to 4 × 4 mm), accommodating diverse applications from small biopsies to larger anatomical regions. When integrated with adjacent-section complementary modalities (such as spatial ATAC-seq^28^), it enables comprehensive interrogation of the gene regulatory logic, spanning chromatin conformation, transcription factor binding, chromatin accessibility, and transcription within a unified spatial context. Moreover, strategic design in our study including histology-guided sampling in melanoma and high-resolution profiling of embryonic tissues, maximized biological insight while maintaining technical feasibility, illustrating principles applicable to future spatial multi-omics efforts.

Together, these findings establish Spatial Hi-C-RNA as a robust and versatile platform for mapping chromatin–gene coupling within intact tissues. By integrating chromatin architecture with transcription at high spatial resolution, Spatial Hi-C-RNA provides direct insight into how 3D genome organization shapes cell identity, developmental trajectories, and tumor progression within native tissue microenvironments. This framework opens new avenues for understanding the structural basis of gene regulation in both physiological and pathological contexts and lays the groundwork for future efforts to integrate chromatin architecture with additional spatial modalities, computational models, and clinical histopathology.

### Limitations of the study

Despite its broad utility, Spatial Hi-C-RNA has several technical constraints that point toward future development. First, the spatial resolution is physically bound by the dimensions of the microfluidic channels employed for deterministic barcoding, a constraint shared with other microfluidic platforms. Although the present implementation achieves near–single-cell resolution, pushing toward subcellular profiling will require next-generation nanofluidic designs or the integration of advanced optical barcoding strategies to overcome fluidic resolution limits. Second, the transcriptomic modality relies on poly(A)-based capture, which excludes important classes of non-polyadenylated RNAs, including many long noncoding RNAs and histone RNAs. Incorporating random-priming or capture-by-ligation methods in future development will provide a more comprehensive view of the transcriptome linked to 3D genome organization. Third, the current chromatin conformation workflow utilizes restriction enzyme digestion, which introduces sequence-dependent fragmentation bias and limits uniformity of genome coverage. Transitioning to sequence-independent fragmentation approaches will alleviate this bias and improve the resolution and accuracy of chromatin contact maps. Addressing these challenges will further enhance the precision, completeness, and interpretability of spatial multi-omics insights.

## RESOURCE AVAILABILITY

### Data and code availability

Raw and processed sequencing data for this study can be accessed in the NCBI Gene Expression Omnibus (GEO) database under the accession number GSE311199. All original code has been deposited at GitHub https://github.com/PengfeiGuo0123/Spatial-Hi-C-RNA) or Zenodo (DOI: 10.5281/zenodo.21475571).

## ACKNOWLEDGMENTS

We acknowledge support from the National Institute on Aging and the National Institute of Allergy and Infectious Diseases of the National Institutes of Health under award numbers R01AG085344 and DP2AI177913 (to Y.D.), the Packard Fellowship for Science and Engineering (to Y.D.), a grant from the Alzheimer’s Association under award number AARG-NTF-23-1150285 (to Y.D.), the pilot award from the Epigenetics Institute at the University of Pennsylvania (to Y.D.), the National Institute of Mental Health under award number R37MH120269 (to J.E.P-C), the National Institute of Neurological Disorders and Stroke under award number R01NS114226 (to J.E.P-C), National Institutes of Health Common Fund/ National Institute of Mental Health under award number DP1MH129957 (to J.E.P-C), the CZI Neurodegenerative Disease Pairs Award under award number DAF2022-250430 (to J.E.P-C). The content is solely the responsibility of the authors and does not necessarily represent the official views of the National Institutes of Health. Computational analyses were performed using the Betty high-performance computing cluster at the Penn Advanced Research Computing Center.

## AUTHOR CONTRIBUTIONS

Conceptualization, P.G. and Y.D.; Methodology, P.G., A.J.W., J.E.P-C., and Y.D.; Experimental investigation, P.G., J.Z.; Data analysis, P.G., Y.C., J.H., Y.F.C., J.T.Z., and Y.D.; Resources, Z.H.; Writing – original draft, P.G., Y.C., and Y.D.; All authors reviewed, edited, and approved the paper.

## STAR*METHODS

### KEY RESOURCES TABLE

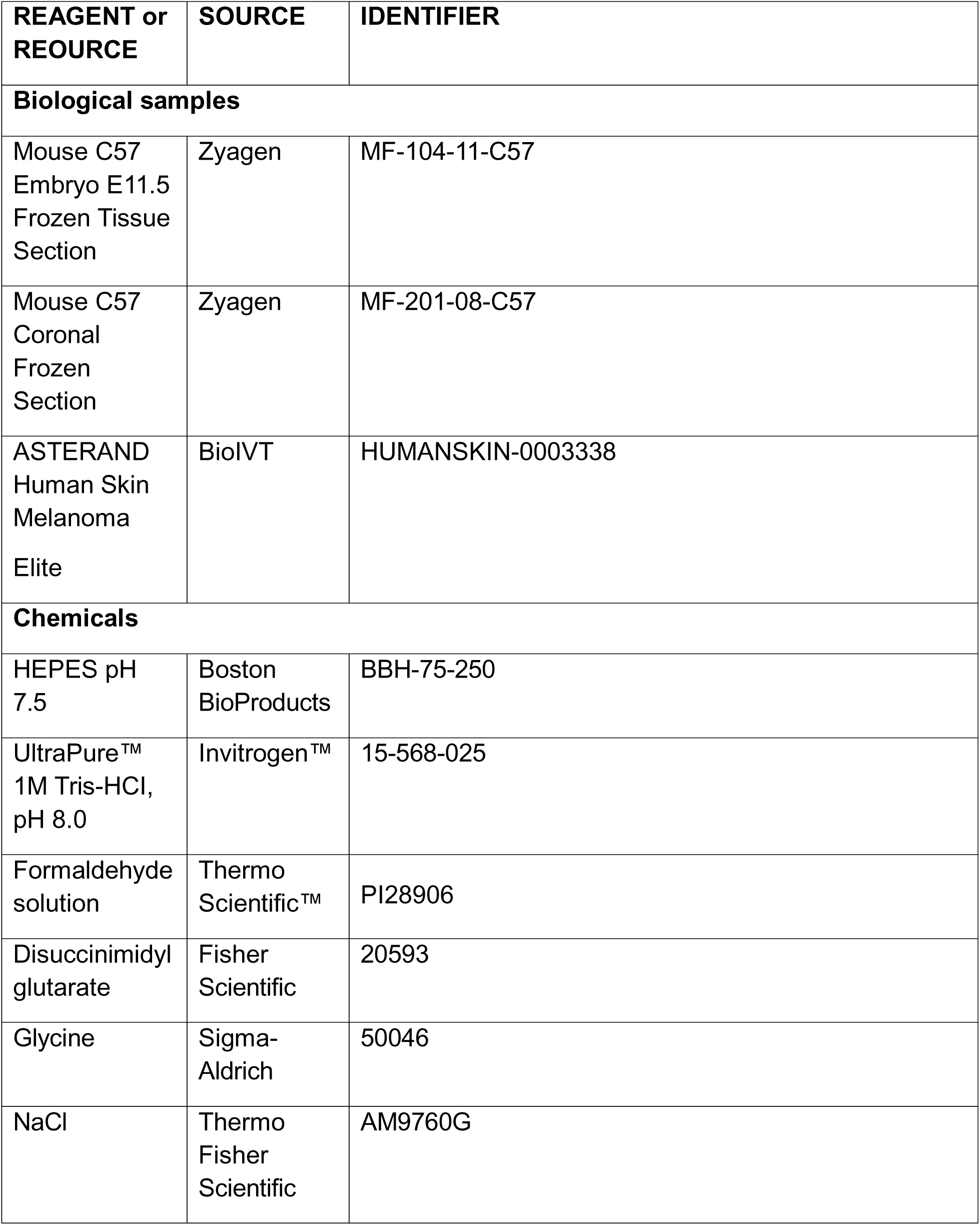
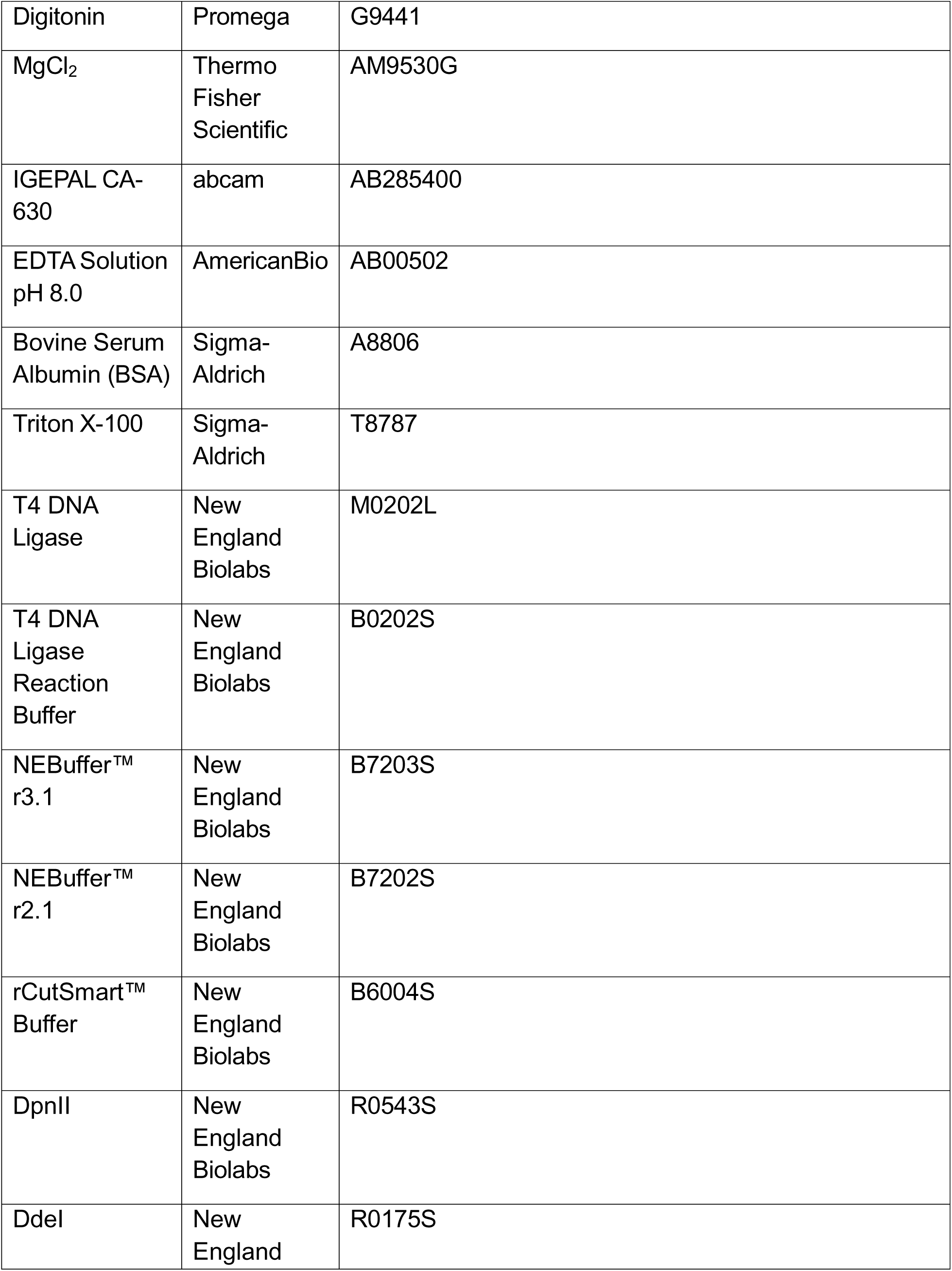
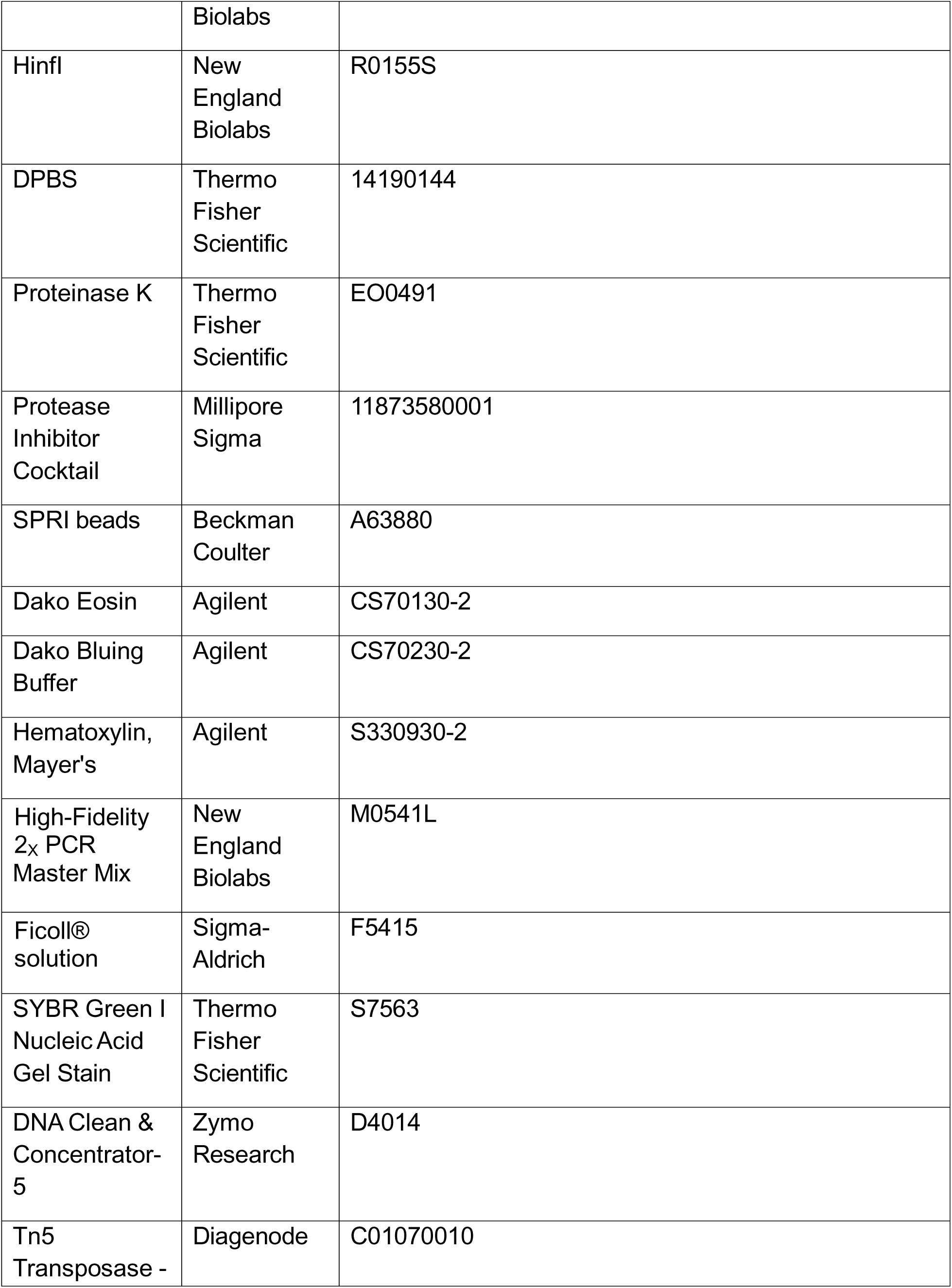
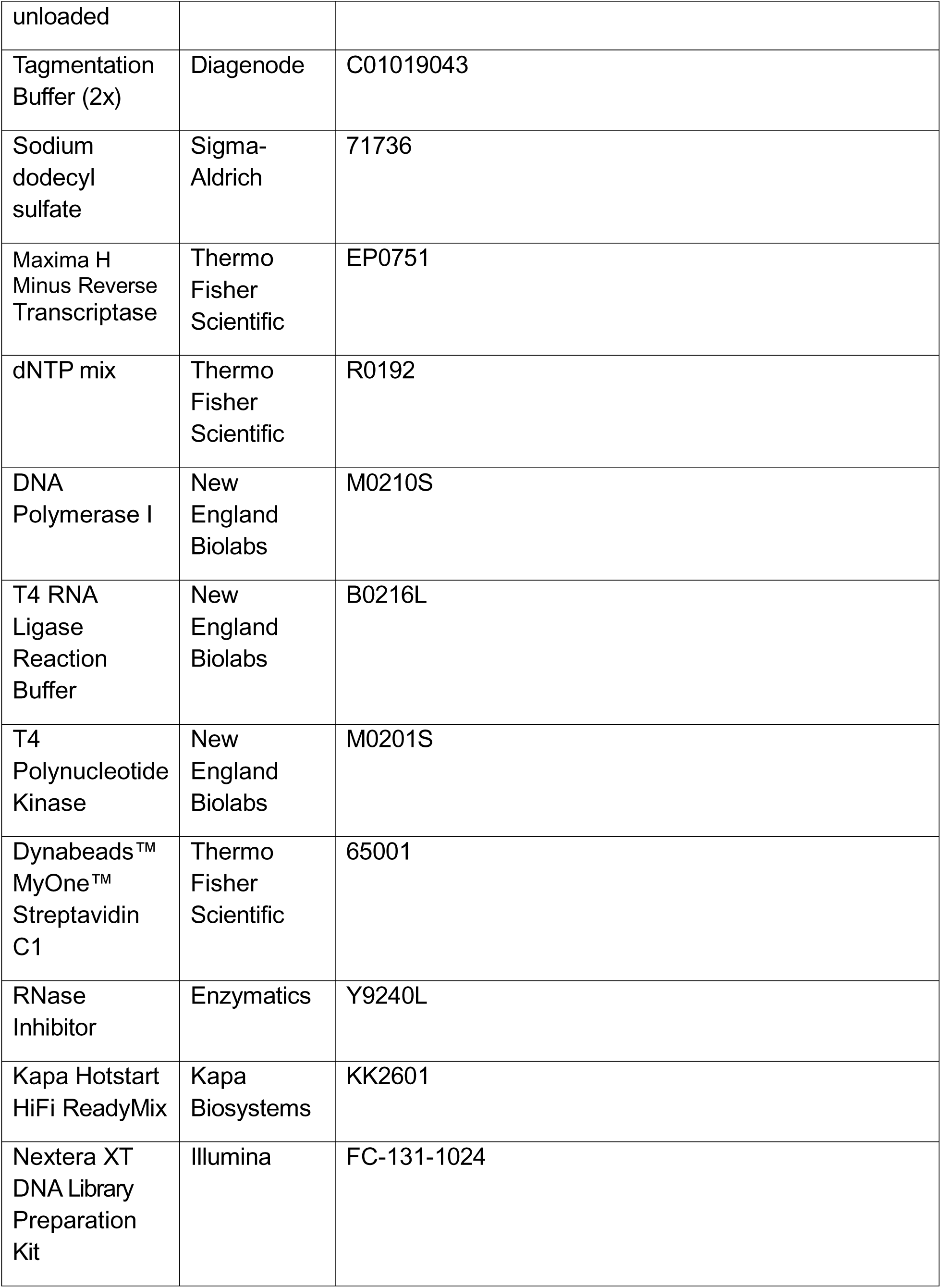
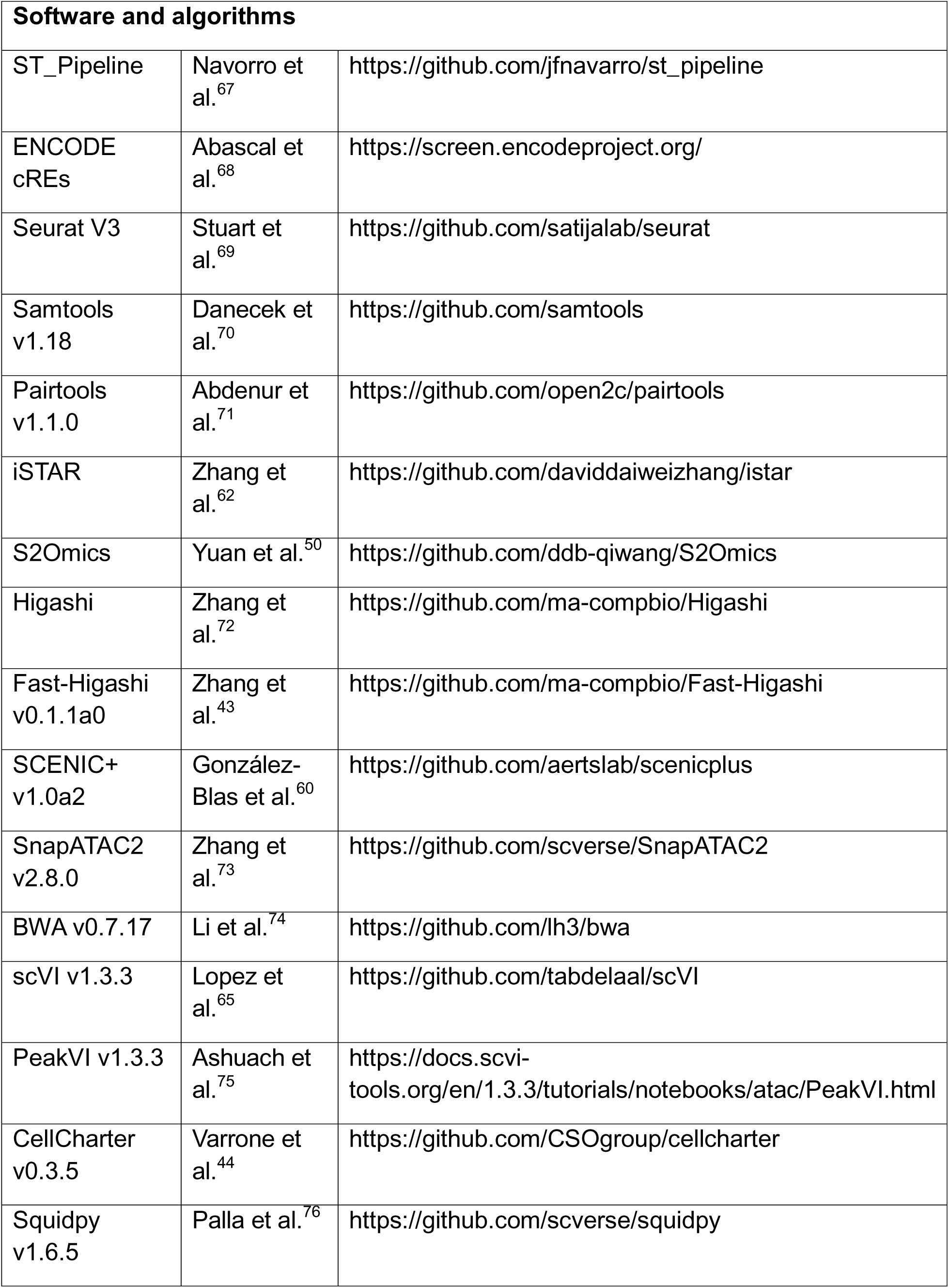
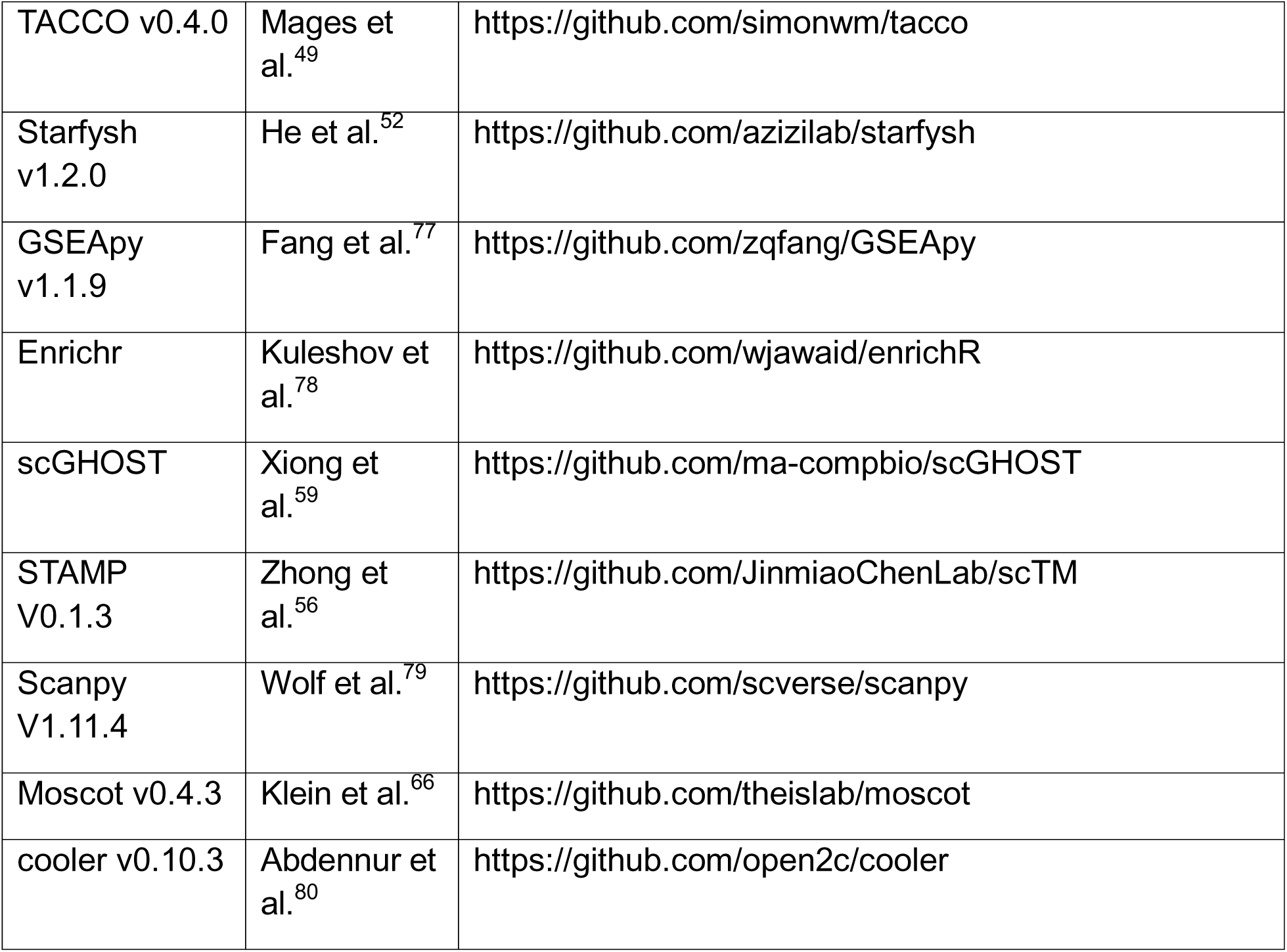

### EXPERIMENTAL MODEL AND STUDY PARTICIPANT DETAILS

#### Sample handling and section preparation

The mouse E11.5 and caudal hippocampus coronal brain sections were purchased from Zyagen (San Diego, CA). Tissues were freshly harvested from C57bl/6 mice. The in-house Institutional Animal Care and Use Committee of Zyagen review and approve all protocols. The human melanoma block was purchased from BioIVT (North America). For both mouse and human samples, tissue sections were mounted on the center of adhesion super frost+ slides (2x 3’’) with a thickness of 10 µm.

### METHOD DETAILS

#### Microfluidic device fabrication and assembly

The molds for polydimethylsiloxane (PDMS) microfluidic devices were fabricated using standard photolithography. The manufacturer’s guidelines were followed to spin-coat SU-8-negative photoresist (nos. SU-2025, Microchem) onto a silicon wafer (no. C04004, WaferPro). The heights of the features were about 20 μm for 20-μm-wide devices, 10 μm for 10-μm-wide devices, respectively. We mixed the curing and base agents in a 1:10 ratio and poured the mixture onto the molds. After degassing for 30 min the mixture was cured at 70°C for 2 h. Solidified PDMS was extracted for further use. The fabrication and preparation of the PDMS device follow the published protocol.^81^

#### DNA barcodes annealing and preparation of the Tn5 transposome

DNA oligos were purchased from IDT (Coralville, IA) and the sequences were listed in Tabel S5. For spatial barcodes, barcode (100 µm) and ligation linker (100 µm) were annealed at 1:1 ratio in 2x annealing buffer (100 mM NaCl, 20 mM Tris-HCl pH 8.0, 2 mM EDTA) with the PCR program (95°C for 5 min, cooling to 20°C at a rate of 0.1°C/s, followed by 12°C for 3 min. Unloaded Tn5 transposase (C01070010) was purchased from Diagenode, and the transposome was assembled according to the manufacturer’s guidelines. The transposome was assembled by combination of Tn5MErev and Tn5ME- A or Tn5ME-B. For Hi-C, Tn5 was assembled with Tn5MErev/ME-B only. For ATAC-seq, Tn5 was assembled with Tn5MErev/ME-B and Tn5MErev/ME-A.

#### Tissue H&E staining

The adjacent tissue section was performed with tissue histology examination using H&E staining. The tissue slide was first fixed with 4% formaldehyde and then stained with the hematoxylin (Agilent) for one minute. Afterward, the slides were incubated in a bluing reagent (Agilent) for 45 seconds. Finally, the slides were stained with eosin (Agilent) for one minute.

#### Spatial co-profiling of 3D chromatin architecture and gene expression Fixation and permeabilization

The frozen tissue section was first thawed for 1 min at 37°C. The tissue was then fixed with 1% formaldehyde plus 0.05 U/µL RNase inhibitor (QIAGEN) for 5 min and quenched with 1.25 M glycine for 5 min. The tissue was re-fixed with 3 mM DSG^41^ plus 0.05 U/µL RNase inhibitor for 40 min and quenched with 1.25 M glycine for 5 min. After fixation, the tissue was washed with deionized H_2_O. Intact tissue morphology was scanned using a 10x objective on the KEYENCE BZ-X800 microscope.

The tissue was subsequently permeabilized with NP-40 lysis buffer (0.2% IGEPAL CA- 630; 10 mM Tris-HCl, pH 8.0; 10 mM NaCl; 1x protease inhibitor; 0.05 U/µL RNase inhibitor) for 15 min, followed by 1x DPBS plus 0.05 U/µL RNase inhibitor to stop permeabilization. The tissue was incubated with 0.5% SDS at 65°C for 10 min and quenched with 1% Triton X-100 plus 0.05 U/µL RNase inhibitor for 5 min. After quenching, the tissue was washed with 0.5x DPBS plus 0.05 U/µL RNase inhibitor for 5 min.

#### In situ reverse transcript

For the in situ reverse transcription, add the RT mixture (12.5 µL 5x reverse transcription buffer, 6.2 µL Maxima H Minus Reverse Transcriptase, 20.3 µL 0.5x DPBS, 9.6 µL RNase free water, 0.8 µL RNase Inhibitor, 10 µL RT primer (100 µM)) and incubate for 30 min at room temperature, then at 42°C for 90 min in a humidified container. After the RT reaction, wash the tissue with 1x NEBr3.1 buffer plus 0.05 U/µL RNase inhibitor for 5 min.

#### In situ Hi-C

The Spatial Hi-C was adapted from published in situ Hi-C protocol.^41,58^ The tissue section was digested with 200 µL restriction enzyme mix (1x NEBr3.1 buffer, 1 U/µL DpnII, 1 U/µL DdeI, 1 U/µL HinfI, 0.05 U/µL RNase inhibitor, 0.1% Triton X-100) and incubated at 37°C overnight. The tissue was then washed with 1x NEBr2.1 buffer plus 0.05 U/µL RNase inhibitor for 5 min. Next, the following mixture (1x NEBr2.1, 100 µM dNTP mix, 0.2 U/µL DNA Polymerase I) was added to blunt the DNA ends and incubated at room temperature for 2 h. Afterwards, wash the tissue with 1x T4 DNA ligase buffer plus 0.05 U/µL RNase inhibitor for 5 min. The proximity ligation mixture (1x T4 DNA ligase buffer, 16 U/µL T4 DNA ligase, 0.1% Triton X-100, 0.05 U/µL RNase inhibitor) was added to the tissue section and incubated at room temperature overnight. Next, the tissue section was further treated with 0.1% SDS and incubated at 55°C for 1 h, then quenched with 1% Triton X-100 plus 0.05 U/µL RNase inhibitor for 5 min.

The tissue section was then washed twice with wash buffer (10 mM Tris-HCl pH 7.4, 10 mM NaCl, 3 mM MgCl_2_, 1% BSA, 0.1% Tween 20, 0.05 U/µL RNase inhibitor) for 5 min each time. Next, 50 µL Tn5 transposition mixture (1.87 μL RNase free water 5 µL Tn5MErev/ME-B transposome, 16.5 µL 1x DPBS, 25 µL 2x Tagmentation buffer, 0.5 µL 1% digitonin, 0.5 µL 10% Tween 20, 0.05 U/µL RNase inhibitor) was added and incubated at 55°C for 1 h, then quenched with 40 mM EDTA plus 0.05 U/µL RNase inhibitor for 5 min. Next, the tissue section was washed twice with DPBS plus 0.05 U/µL RNase inhibitor for 5 min and then proceeding for in situ barcoding.

#### In situ barcoding and DAPI staining

For the first ligation of barcode Ai, the PDMS chip A was attached to the tissue region of interest (ROI), then the 10x objective (KEYENCE BZ-X800 microscope) was performed to capture the position of the aligned chipA-tissue image. Next, an acrylic clamp was applied to tightly fix the PDMS to the tissue slide. Ligation master mixture was prepared for each inlet channel with 1.5 µL annealed DNA barcode Ai (i = 80 or 200) and 4 µL ligation mixture (e.g. for 100 channels: 288.2 µL RNase-free water, 23.2 µL NEB buffer r3.1, 54 µL T4 DNA ligase buffer, 1.8 µL RNase inhibitor, 10.8 µL 5% Triton X-100, 22 µL T4 DNA ligase). For each inlet, load 4.5 µL ligation master mixture and apply the vacuum to flow the mixture over the ROI of tissue section, then incubate at 37°C for 30 min in a humidified container. After incubation, remove the PDMS chip and clam to wash the tissue section with deionized water. For the second ligation of barcode Bj, the PDMS chip B was covered to the ROI and the bright-field imaged was captured. Subsequently, the ligation of DNA barcode Bj was performed similarly to barcode Ai set. Finally, the tissue section was washed with 1x DPBS and dried for DAPI staining. The tissue section was incubated with 300 nM DAPI solution at room temperature for 10 min. After twice washes with RNase-free water, the final scan was performed with the 10x objective to scan the DAPI staining and bright-field image of the tissue section.

#### Tissue lysis and gDNA and cDNA library separation

The tissue ROI was covered with a PDMS reservoir and digested with 100 µL lysis mixture (0.4 mg/mL proteinase K, 1 mM EDTA, 50 mM Tris-HCl pH 8.0, 200 mM NaCl, 1% SDS) at 58°C for 2 h in a wet box. The lysate was then collected in a 0.2 ml tube and incubated at 60°C overnight. For gDNA (Hi-C) and cDNA separation, the lysate first was purified with Zymo DNA Clean & Concentrator-5 column and eluted with 100 µL of RNase-free water. The lysate was mixed with 40 µL of Dynabeads MyOne Streptavidin C1 beads resuspended in 100 µL of 2x B&W buffer (10 mM Tris-HCl pH 7.5, 1 mM EDTA, 2 M NaCl, 0.05 U/µL RNase inhibitor) and the mixture was incubated at room temperature for 1 h with agitation. A magnetic rack was used to separate beads (containing cDNA) and supernatant (containing gDNA) in the eluent. The supernatant was collected and purified with with Zymo DNA Clean & Concentrator-5 again and eluted with 20 µL of RNase-free water for Hi-C library construction.

#### Hi-C library generation

The eluted gDNA (Hi-C) was pre-heated at 96°C for 5 min and then cooled down on ice for 2 min. Prepare 12 µM splint ligation P5 adapter mixture (30.6 µL H20, 6 µL SL_P5_rc primer (100 µM), 8.4 µL S5MEA_N10 primer (100 µM), 5 µL 10x T4 RNA ligase buffer) with the following program: 95°C for 1 min, cooling to 10°C at a rate of 0.1°C/s, followed by 10°C for 5 min. This adapter mixture was diluted to 0.75 µM and 10 µL adapter mixture was mixed with gDNA, followed by adding 80 µL ligation master mixture (40 µL 50% PEG-8000, 12.5 µL SCR buffer (666 mM Tris-HCl, pH 8.0 and 132 mM MgCl_2_), 10 µL 100 mM DTT, 10 mM ATP, 1.25 µL T4 Polynucleotide Kinase (10 U/µL), 6.25 µL T4 ligase (400 U/µL)). Next, the mixture was incubated with the following program: 37°C for 45 min, 65°C for 20 min. Afterwards, 200 µL PCR mixture (150 µL High-Fidelity 2x PCR Master Mix, 5 µL N70X (10 µM), 5 µL N50X (10 µM), 40 µL H_2_O) was added to gDNA mixture, and run with the following PCR program: 95°C for 5 min, then cycling at 95°C for 30 s, 56°C for 30 s and 72°C for 60 s, for 10–18 cycles, followed by 72°C for 5 min. The amplified PCR product was purified with 0.7x SPRI beads, and the Hi-C library was eluted with 20 µL H_2_O.

#### RNA library generation

The separated beads were used for cDNA library construction. They were washed twice with 400 µL of 1× B&W buffer (containing 0.05% Tween-20 containing and 0.05 U/µL RNase inhibitor) and once with 10 mM Tris pH 8.0 (containing 0.1% Tween-20 and 0.05 U/µL RNase inhibitor). The separated beads were further washed with 400 µL RNase-free water. Streptavidin beads with bound cDNA molecules were resuspended in 200 µL TSO mixture (22 µL 10 mM dNTPs, 44 µL 5× Maxima RT buffer, 44 µL 20% Ficoll PM- 400 solution, 88 µL H_2_O, 5.5 µL 100 uM template switch primer, 11 µL Maxima H Minus Reverse Transcriptase, 5.5 µL RNase Inhibitor) and incubated at room temperature for 30 min and then at 42°C for 1.5 h. After incubation, beads were washed once with 400 µL 10 mM Tris and 0.1% Tween-20 and then with RNase-free water. Washed beads were resuspended in 220 µL PCR mixture (110 µL 2× Kapa HiFi HotStart Master Mix, 8.8 µL 10 μM PCR primer 1 and primer 2, 92.4 µL H_2_O), then split 50 µL beads mixture per tube, and run on PCR thermocycling with the following program: 95°C for 3 min, cycling at 98°C for 20 s, 65°C for 45 s and 72°C for 3 min with 5 cycles. After PCR reactions, the product was separated from beads and SYBR Green was added at a final 1x concentration. The PCR product was amplified on a qPCR machine with the following conditions: 95°C for 3 min, cycling at 98°C for 20 s, 65°C for 20 s and 72°C for 3 min, 15 times, followed by 5 min at 72°C. The reaction was stopped once the curve signal reached the plateau. The PCR product was then purified with 0.6x SPRI beads and eluted in 20 µL RNase-free. Next, the Nextera XT DNA Library Prep Kit was used for the RNA library generation. In brief, 2 ng purified PCR product was diluted in RNase-free water to a total volume of 5 µL, then 10 µL Tagment DNA buffer and 5 µL Amplicon Tagment mix were added and incubated at 55°C for 5 min; 5 µL NT buffer was added to stop the tagmentation at room temperature for 5 min. 25 μl µL PCR master mixture (15 µL PCR master mix, 1 µL N50X primer and 1 μL N70X primer (10 μM), 8 μL H_2_O) was added to the tagmented DNA product and run with the following program: 95°C for 30 s, cycling at 95°C for 10 s, 55°C for 30 s, 72°C for 30 s and 72°C for 5 min, for 12 cycles. The PCR product was purified with 0.7x SPRI beads to obtain the final RNA library.

#### Library QC and sequencing

The Agilent D5000 Screentape was used to determine the size distribution and concentration of the library before sequencing. NGS was conducted on an Illumina NovaSeq X Plus sequencer (paired-end, 150-base-pair mode).

#### Spatial profiling of chromatin accessibility

Spatial ATAC-seq was performed with frozen tissue section. Briefly, the tissue section underwent fixation, permeabilization, and tagmentation, followed by spatial barcoding as described above. The library construction followed the protocol as described in 2022.^28^

#### Processing of Spatial Hi-C-RNA and spatial ATAC-seq raw data

For the ATAC-seq data, linkers 1 and 2 were used to selectively filter Read 2 before alignment with BWA v0.7.17,^74^ after which the resulting files were sorted and indexed using Samtools v1.18^70^ to streamline data processing. During this step, genomic sequences were assigned to Read 1, while barcodes A and B were incorporated into Read 2. The resulting FASTQ files were mapped to the human reference genome (GRCh38). This workflow generated fragment files in a TSV-like format, in which each entry contains both genomic coordinates and the corresponding spatial barcode pair, enabling integrated downstream analyses.

For the Hi-C data, linkers 1 and 2 were used to selectively filter Read 2 before alignment. For alignment, BWA-MEM V0.7.17,^74^ was used with specified arguments ‘-SP5M’. After alignment, bam file was used as input for parse, sort, and deduplicate by Pairtools V1.1.0.^71^ The barcode information was stored in a separate column in the pair file. Contact matrices in the .mcool format were generated and balanced using the cooler v0.10.3.^80^

For the RNA-sequencing data, read 2 was parsed to retrieve barcode A, barcode B, and the UMI. The resulting reads were then processed with the ST Pipeline^67^ and aligned to the mouse reference genome (GRCm38 or GRCh38). This workflow generated a gene-by-barcode matrix in which each expression profile is linked to its spatial location through the paired barcodes A and B, providing a foundation for spatially resolved transcriptomic analyses.

#### Analysis of Spatial Hi-C-RNA and spatial ATAC-seq data

##### Spatial Hi-C contact frequency

For each spatially barcoded Hi-C dataset, intra-chromosomal contacts were extracted and genomic distances between paired anchors computed. Distances were binned on a log₂ scale (step size 0.125), ranging from 10 kb to 195 Mb for mouse and 10 kb to 250 Mb for human samples. The proportion of contacts within short-range (100 kb–2 Mb) and long-range (20–100 Mb) bins was calculated for each barcode, and a short/long ratio was defined as the quotient of these proportions. These ratios were merged with spatial coordinates, log-transformed, and visualized as a spatial scatter plot to map distance-dependent chromatin contact decay across the tissue.

##### Spatial embedding, clustering, and visualization

Single-modality spatial clusters were generated using the standard CellCharter v0.3.5^44^ workflow. Spot or cell features were first embedded using the user-selected method and then spatially aggregated to produce spatial representations, which were clustered using a Gaussian mixture model (GMM). For multimodal clustering, feature representations from each modality were concatenated before spatial aggregation, and the same GMM-based procedure was applied.

##### RNA modality embedding

For the RNA modality, per-spot representations were obtained using scVI v1.3.3.^65^ RNA data were first filtered with scanpy.pp.filter_cells() and scanpy.pp.filter_genes() (min_genes = 10; min_cells = 10) to remove low-quality cells and lowly expressed genes. Highly variable genes were selected using scanpy.pp.highly_variable_genes() with the Seurat_V3 method (5,000 genes for mouse brain, 2,000 for human melanoma, and 4,000 for mouse embryo). Raw counts were then used to train scVI with gene_likelihood=’nb’, and both n_latent and n_hidden set to 256, with early stopping enabled. Latent representations for each spot were extracted from the pretrained model using get_latent_representation() function.

##### Hi-C modality embedding

For the Hi-C modality, per-spot representations were generated using Fast-Higashi v0.1.1a0.^43^ Training was performed at 500-kb resolution with parameters dim1 = 0.6, rank = 256, and n_iter_parafac = 1. Fast-Higashi includes an internal quality-control module that restricts embedding to high-quality spots or cells and propagates learned representations to lower-quality inputs, eliminating the need for additional preprocessing. Final Hi-C embeddings were obtained as the L2-normalized output of the fetch_cell_embedding() function.

##### ATAC modality embedding

For the ATAC modality, preprocessing was performed using SnapATAC2 v2.8.0^73^ to call peaks from fragment files based on the human ENCODE cREs^68^ reference. A peak-by-spot matrix was generated with snapatac2.pp.make_peak_matrix() in paired-insertion mode, and highly variable peaks were selected using scanpy.pp.highly_variable_genes() with n_top_genes = 250,000. The resulting matrix was used to train PeakVI v1.3.3^75^ with n_latent = 16, and per-spot ATAC representations were obtained using get_latent_representation() function.

##### Spatial clustering

For single-modality spatial clustering of RNA, Hi-C, and ATAC data, modality-specific representations were used as input to cellcharter.gr.aggregate_neighbors(). Spatial neighborhood graphs were first constructed with Squidpy v1.6.5^76^ using squidpy.gr.spatial_neighbors() in grid mode (n_neigh = 8), consistent with the grid-based layout of the dataset. Spatial embeddings were then generated with cellcharter.gr.aggregate_neighbors(), using n_layers = 2 for mouse embryo, mouse brain, and melanoma RNA, Hi-C, and integrated data, and n_layers = 3 for melanoma ATAC.

Clustering was performed with the Gaussian mixture model implemented in cellcharter.tl.Cluster(), and the optimal number of clusters was selected based on the best convergence across repeated runs (random seed = 42, GPU mode enabled). This procedure assigned a spatial cluster label to each spot. For integrated clustering, multimodal representations were obtained by concatenating RNA and Hi-C embeddings to retain modality-specific structure without imposing parametric coupling. The same spatial aggregation and clustering steps used in the single-modality analysis were applied to generate multimodal spatial clusters. For the mouse embryo dataset, clusters containing fewer than 10 cells (n_cells < 10) were removed to obtain the final cluster assignments.

##### Cell type deconvolution, annotation transfer, and downstream analysis

All analyses in this section used the RNA modality as input for cell type deconvolution or label transfer. The mouse embryo dataset (10 μm resolution) was analyzed using label transfer, whereas the mouse brain and human melanoma datasets (20 μm resolution) were processed using cell type deconvolution. For mouse brain and embryo, cell type annotation was performed using TACCO v0.4.0.^49^ Raw counts from the spatial data and reference atlases were provided as input to tacco.tl.annotation, which produced a spot-by–cell type proportion matrix. Spots with a dominant cell type (i.e., the highest proportion exceeding a predefined threshold) were assigned a cell type label for downstream analyses. For human melanoma, cell-state deconvolution was performed using Starfysh v1.2.0.^52^ Curated marker sets representing major immune, fibroblast, and melanoma states were provided, and raw RNA counts were processed through the official Starfysh workflow. The resulting spot-by–cell type proportion matrix was treated analogously to the TACCO output, with spots dominated by a single state assigned the corresponding cell type label.

##### Cell–cell interaction difference

CellCharter.gr.enrichment() was applied using spatial cluster labels (from RNA, Hi-C, and integrated modalities) to define groups and cell types to define categories. This approach compares each cluster’s cell type composition to the global background to calculate fold changes, with higher values indicating stronger enrichment. For melanoma, a maximum-proportion threshold of 0.2 was applied to filter mixed spots and assign final cell type labels.

##### Cell type co-occurrence

Fine-grained spatial relationships beyond immediate neighbors were analyzed using TACCO’s tacco.tl.co_occurrence() with delta_distance = 10 and max_distance = 40. This function calculates a distance-dependent co-occurrence score, representing the normalized likelihood of each cell type occurring near a given central cell type as distance increases, thereby capturing spatial co-localization patterns across multiple scales.

##### Gene set enrichment

Gene set enrichment was performed using the gseapy.enrichr() function from GSEApy v1.1.9^77^ via the Enrichr^78^ API. Significant terms were selected based on an FDR- adjusted p-value threshold specified by the function’s cutoff parameter.

##### Hi-C–specific downstream analysis

###### Higashi model training and imputation

The Higashi v0.1.0a0^72^ was used for A/B compartment scoring, insulation score calculation, TAD identification, and chromosome subcompartment analysis. Due to the sparsity of raw Hi-C data, the Higashi model was first trained (500 kb resolution for A/B compartments, 100 kb for insulation and TADs) using Higashi. train_for_imputation_with_nbr(). Neighborhood-enhanced imputation (Higashi.impute_with_nbr()) generated per-cell Hi-C profiles for downstream analysis, and embeddings for scGHOST-based subcompartment identification were extracted via Higashi.train_for_embeddings().

###### A/B compartment score calculation

Neighborhood-imputed Hi-C profiles were used to calculate A/B compartment scores with the scCompartment scripts from the Higashi repository. CpG density was first computed using the provided script and supplied via the --calib-file parameter to standardize principal component orientation across cells. The output was a cell-by-bin matrix of raw and z-normalized A/B scores, with z-normalized scores used for downstream analyses unless otherwise noted.

###### Insulation score and TAD identification

Neighborhood-imputed Hi-C profiles at 100 kb resolution were analyzed using the scTAD script from the Higashi repository, with window_ins = 100 kb for insulation score calculation and window_tad = 100 kb for TAD identification. The output was a cell-by-bin matrix containing insulation scores and TAD presence annotations (0 = present, 1 = absent).

###### Marker bin selection

Condition-associated bins were identified using a random forest–based feature selection approach. A sklearn.ensemble.RandomForestClassifier() model was trained with condition labels as targets and per-spot Hi-C features (insulation scores, A/B compartment scores, and subcompartment labels) as inputs. Feature importance values were extracted via model.feature_importances_, and the top-ranked bins were selected using a Top K threshold.

###### Subcompartment identification

Chromosome subcompartments were extracted at the spot level using scGHOST^59^ with Higashi-imputed Hi-C profiles, embeddings, and A/B compartment scores as input. The pipeline was run at 500 kb resolution with default parameters, generating a cell-by-bin matrix where each entry represents the bin-level subcompartment (scA1, scA2, scB1, scB2, scB3) under the default five-cluster configuration.

###### Spatial topic decomposition

STAMP V0.1.3^56^ was applied to the RNA modality of the human melanoma dataset to infer spatial transcriptomic topics. Input RNA data were preprocessed to select 2000 highly variable genes using the Seurat_v3 method, and raw counts were retained. STAMP was trained with a learning rate of 0.01, a negative binomial likelihood, and 7 topics. The output is a cell-by-topic proportion matrix, with each topic represented as a weighted combination of genes. Top contributing genes for each topic were identified and subjected to gene set enrichment analysis using gseapy.enrichr() with KEGG Human 2021^82^ gene sets.

###### Inferring spatial tissue structures using iStar

iStar^62^ was used to infer single-cell–resolution spatial Hi-C features. The official iStar implementation was adapted for the microfluidic square-pixel format, instead of its original circular Visium design. Adjacent H&E images, manually registered, provided morphological guidance for super-resolution. Per-cell A/B compartment scores served as the target feature, and the pipeline was executed with a raw pixel size of 0.44 to match the H&E image resolution.

###### Integration with reference datasets and spatiotemporal projection

Cross-dataset integration was performed to obtain a batch-corrected latent space. The RNA modality was jointly integrated with a reference Stereo-seq dataset^66^ using scVI. Both datasets were provided as raw counts, and scVI was run with batch_key specifying the dataset origin and early_stopping=True; all other parameters were kept at default. Per-cell embeddings were extracted using model.get_latent_representation(). Hi-C spatial cluster annotations were then transferred to the Stereo-seq dataset via K-nearest neighbor classification (sklearn.neighbors.KNeighborsClassifier, K = 5). Spatiotemporal mapping of Stereo-seq slices at E10.5 and E11.5 was performed using Moscot v0.4.3^66^ SpatioTemporalProblem, which leverages optimal transport to incorporate spatial structure. Following the official tutorial, the same settings were applied, and the pull and push functions were used to propagate Hi-C cluster information across developmental stages, yielding a matching matrix between source and target cells over time.

###### Pseudotime analysis

Pseudotime inference was performed using Scanpy V1.11.4^79^ diffusion maps (scanpy.tl.diffmap()) based on neighborhood graphs constructed with scanpy.pp.neighbors(). To focus on lineage transitions, maximum-proportion filtering was applied using assigned max-proportion cell type annotations, with thresholds set to 0.5 for the melanoma dataset and 0.2 for the mouse embryo dataset. Pseudotime analysis was conducted on filtered cells, further restricted to selected spatial clusters and lineage-relevant subsets. For Hi-C data, diffusion maps were computed from the top 10 principal components of Fast-Higashi embeddings, while RNA diffusion maps were calculated directly from scVI embeddings. Dimensionality reduction for Hi-C was applied to stabilize variation due to the sparser nature of Fast-Higashi embeddings, in line with previous pipelines.^21^

###### Gene regulatory network analysis

Gene regulatory networks were inferred using adjacent spatial ATAC slices and the RNA modality from co-profiled spatial Hi-C–RNA slices with SCENIC+ v1.0a2.^60^ Spatial ATAC data were manually registered to Hi-C–RNA slices, and tumor/non-tumor regions were labeled by Hi-C clusters. Preprocessed cRE matrices were used for ATAC, and RNA data were filtered and highly variable genes selected (min_genes = 10, min_cells = 10, min_mean = 0.0125, max_mean = 3, min_disp = 0.5). The official SCENIC+ Snakemake workflow was run with is_multiome = False for unpaired multimodal input.

