## Supplementary figures for "Integrative spatial profiling of 3D genome organization and gene expression in tissue"

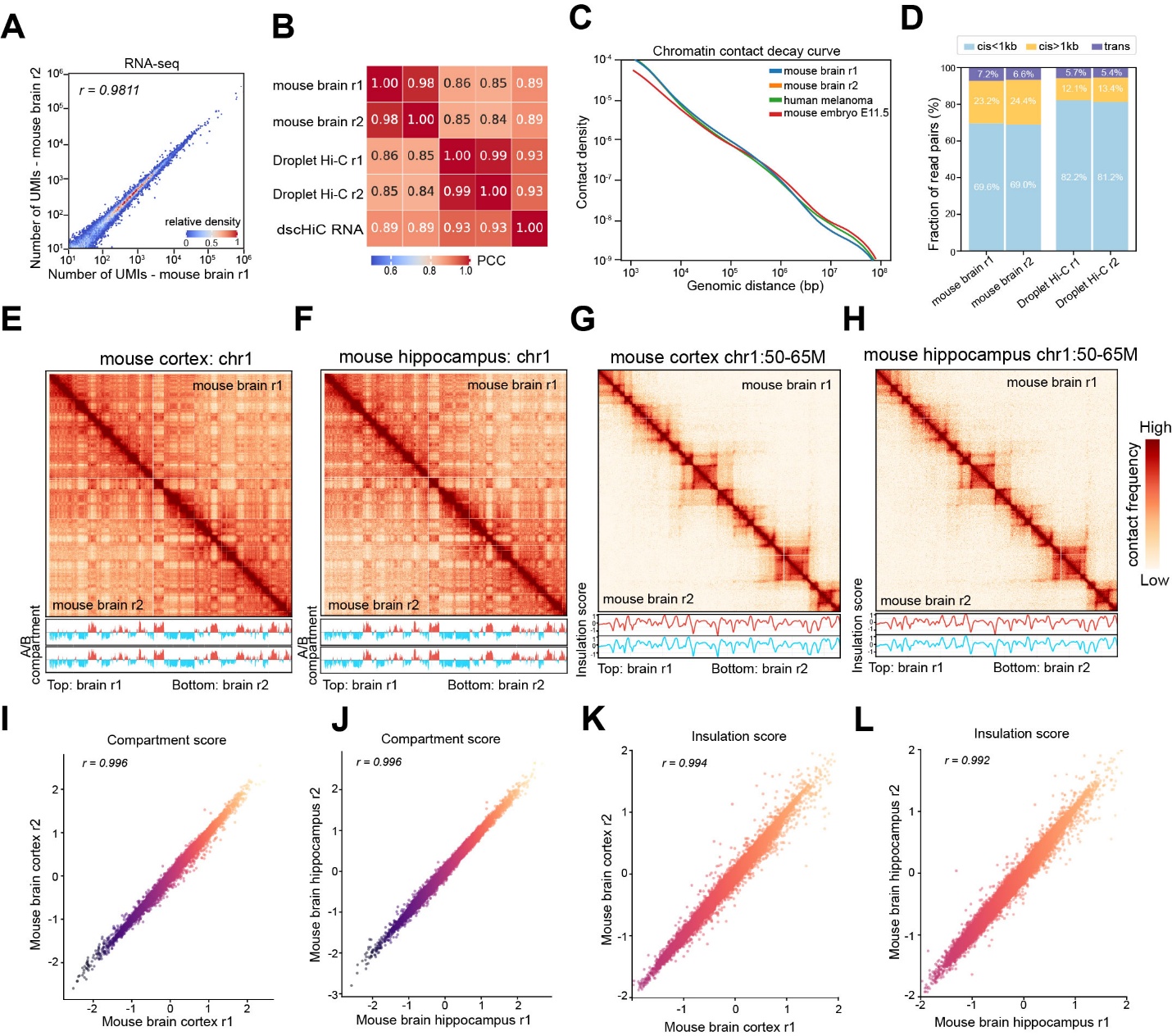

**Figure S1. Spatial Hi-C-RNA performance in the mouse brain sections**

(A) Reproducibility between two biological replicates of RNA libraries from Spatial Hi-C-RNA.

(B) Comparison of aggregated mouse cortex RNA profiles of Spatial Hi-C-RNA replicates with Droplet Hi-C and dscHiC RNA profiles.

(C) Contact frequency by distance across all Spatial Hi-C-RNA datasets.

(D) Comparison of contact ratio in mouse brain datasets generated by Spatial Hi-C and Droplet Hi-C data.

(E-F) Comparison of aggregated contact maps of two Spatial Hi-C replicates from mouse brain cortex (E) and Hippocampus (F) at 100 kb resolution. Bottom tracks are for A/B compartment tracks.

(G-H) Comparison of aggregated contact maps of two Spatial Hi-C replicates from mouse brain cortex (G) and Hippocampus (H) at 25 kb resolution. Bottom tracks are for insulation score tracks.

(I-J) Correlation analysis of compartment values of two Spatial Hi-C replicates from mouse brain cortex (I) and Hippocampus (J).

(K-L) Correlation analysis of insulation scores of two Spatial Hi-C replicates from mouse brain cortex (K) and Hippocampus (L).

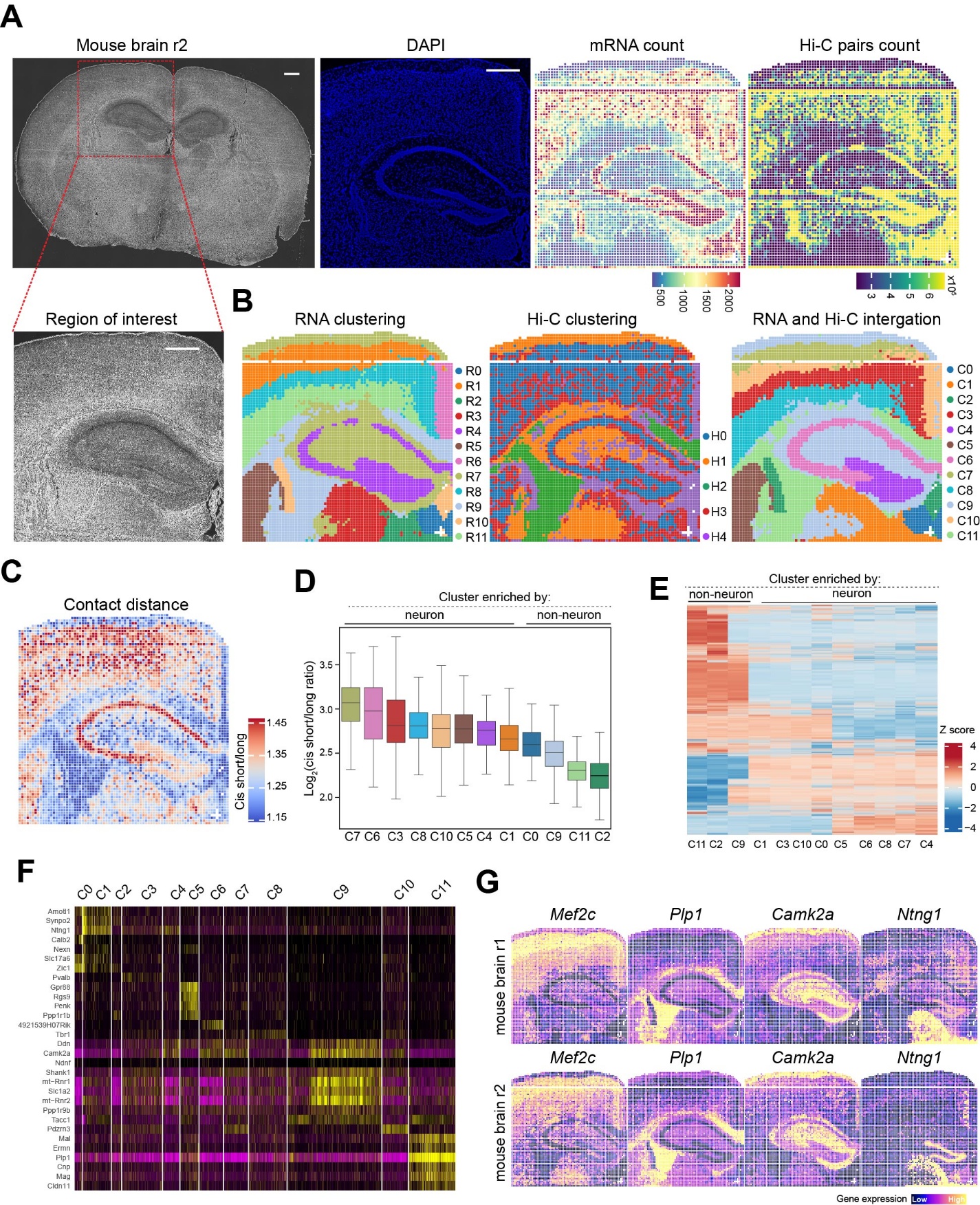

**Figure S2. Spatial Hi-C-RNA performance in the replicate mouse brain section**

(A) Spatial Hi-C-RNA co-profiling of the replicate mouse brain section: overall tissue morphology (left-most), DAPI staining of ROI (left), spatial RNA (right) and Hi-C (right most) count maps. Scale bar: 500 µm.

(B) Spatial distributions of unsupervised RNA (left), Hi-C (middle) and Hi-C/RNA integrated (right) clustering.

(C-D) Spatial pattern of cis short/long contact distance across the mouse brain (C). Distribution of cis short/long ratio in different Hi-C/RNA integrated clusters (D).

(E) Heatmap showing A/B compartment shifts among selected Hi-C clusters, with scaled and sorted compartment scores.

(F) Heatmap showing the normalized gene expression of markers corresponding to each Hi-C/RNA integrated cluster.

(G) Spatial distributions of gene expression for selected marker genes in two mouse brain samples.

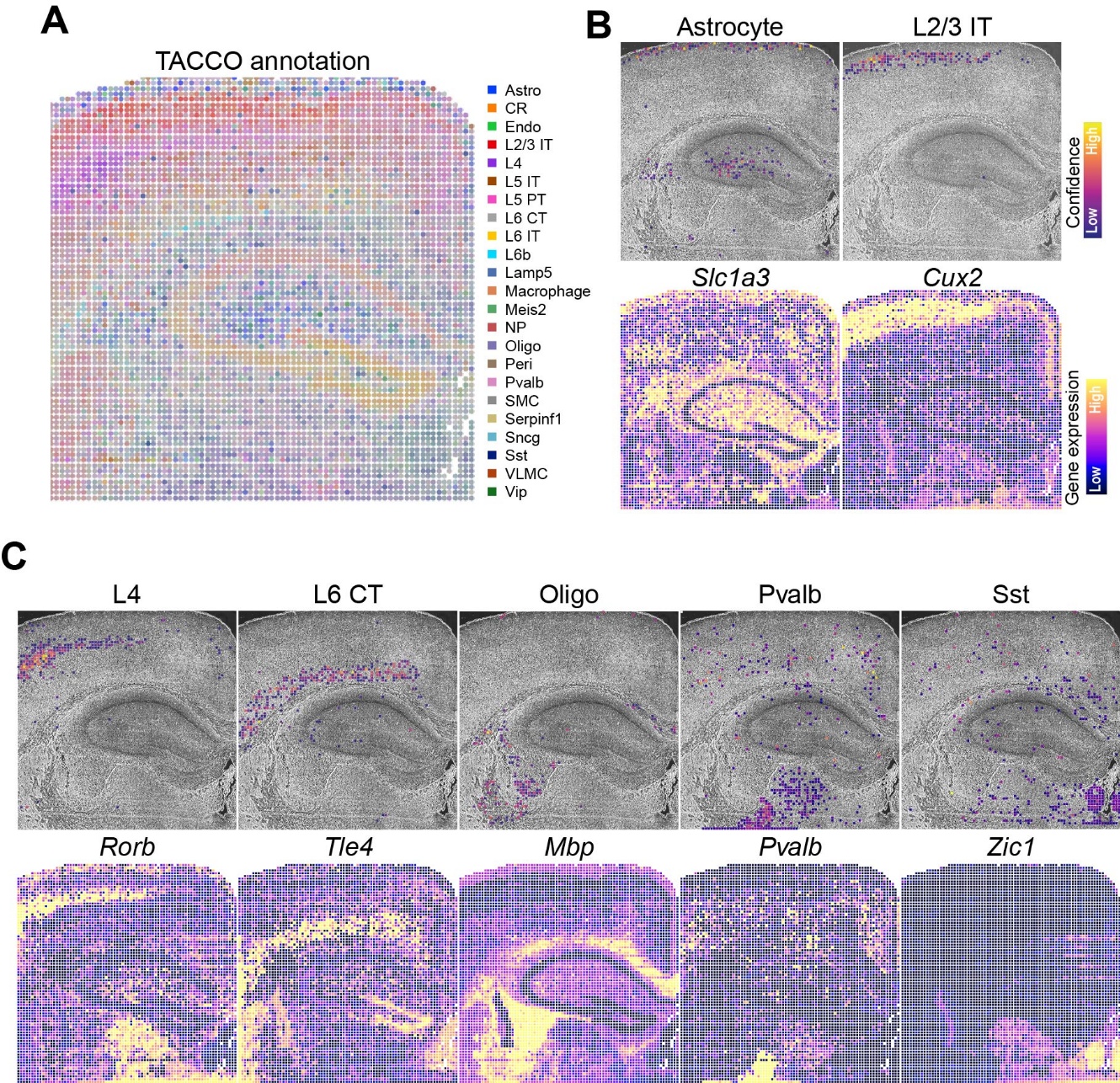

**Figure S3. Spatial cell-type annotation and gene expression mapping in mouse brain**

(A) Cell-type annotation showing the spatial distribution of major cell populations across the mouse brain tissue section.

(B-C) Spatial maps illustrating representative cell-type enrichments (top) and canonical marker gene expression patterns (bottom).

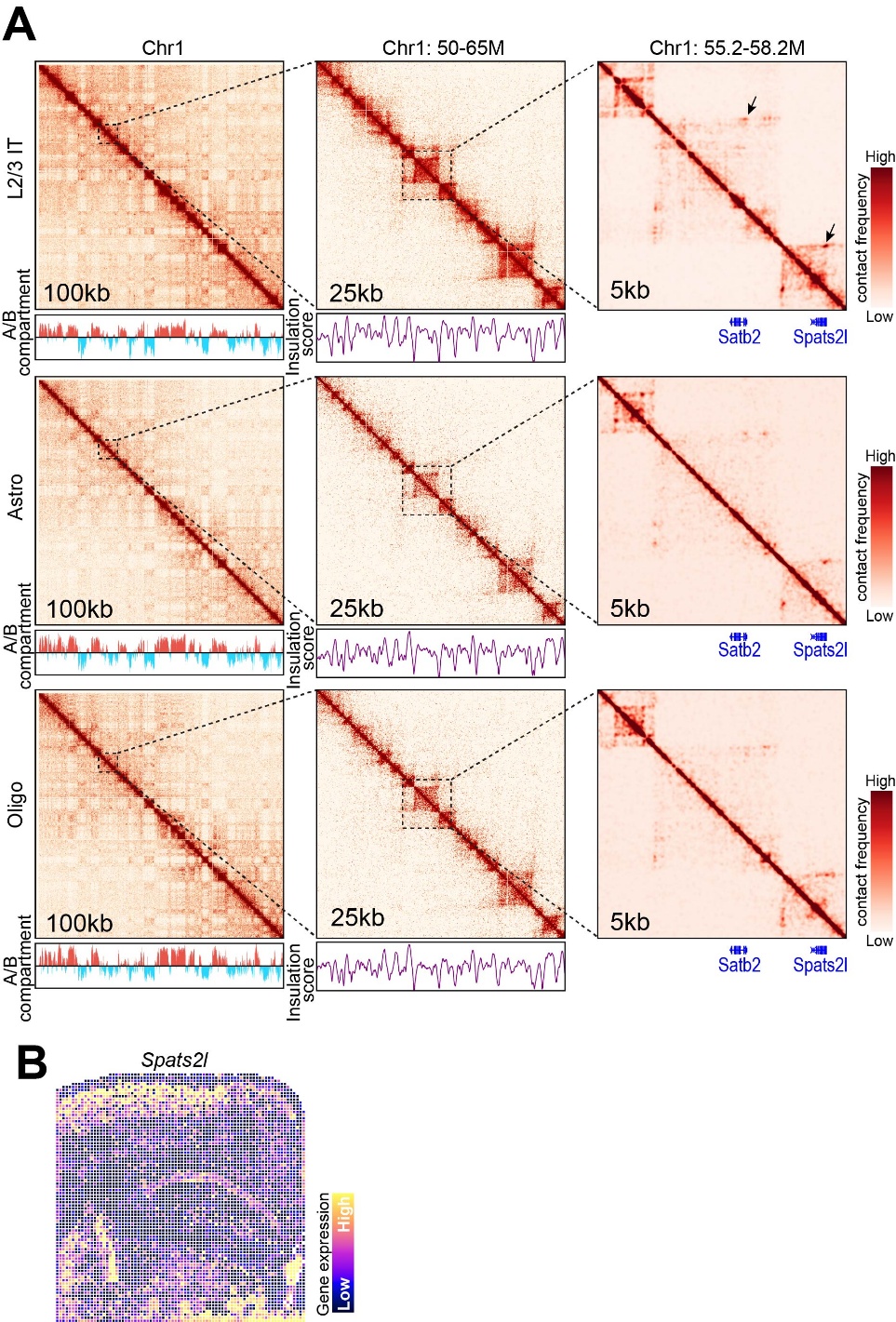

**Figure S4. Cell-type–resolved chromatin architecture reveals fine-scale 3D genome features**

(A) Cell-type–specific Hi-C contact matrices for L2/3 IT neurons, astrocytes, and oligodendrocytes shown at chromosome-wide (100 kb), regional (50–65 Mb, 25 kb), and fine-scale (55.2–58.2 Mb, 5 kb) resolutions. The maps reveal compartments, TAD-like domains, and insulation patterns, with insets highlighting sub-TAD differences and focal loops near the *Satb2–Spats2l* locus that are strongest in L2/3 IT cells and reduced in non-neuron cells.

(B) Spatial gene expression map of *Spats2l* across the tissue section.

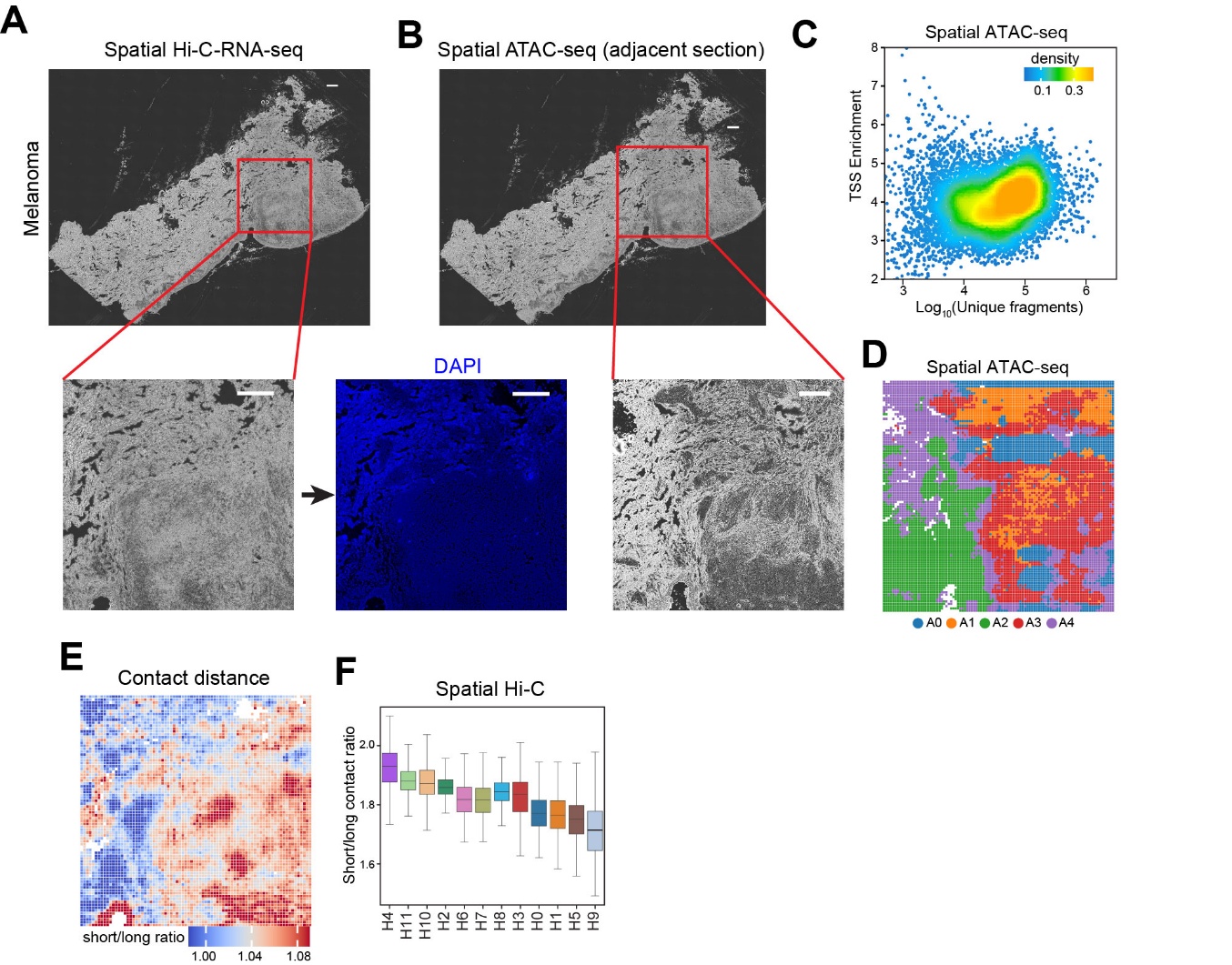

**Figure S5. Spatial multi-omics profiling of melanoma tissue using Spatial Hi-C- RNA and spatial ATAC-seq**

(A) Spatial Hi-C–RNA measurements across a melanoma section (top). The inset highlights the region of interest (bottom left), with the accompanying DAPI image marking nuclear morphology (bottom right). Scale bar: 500 μm.

(B) Spatial ATAC-seq performed on an adjacent section (top). The enlarged inset shows the region of interest (bottom).

(C) Quality assessment of spatial ATAC-seq libraries. Shown is the distribution of TSS enrichment versus log_10_(unique fragments) across spatial pixels.

(D) Spatial ATAC-seq clustering map depicting chromatin accessibility-defined domains (A0–A4), illustrating clear spatial segregation of epigenomic states within the melanoma section.

(E) Contact-distance map derived from spatial Hi-C, showing the ratio of short- to long-range contacts across the tissue.

(F) Distribution of cis short/long contact ratios across Hi-C–defined domains (H0–H11).

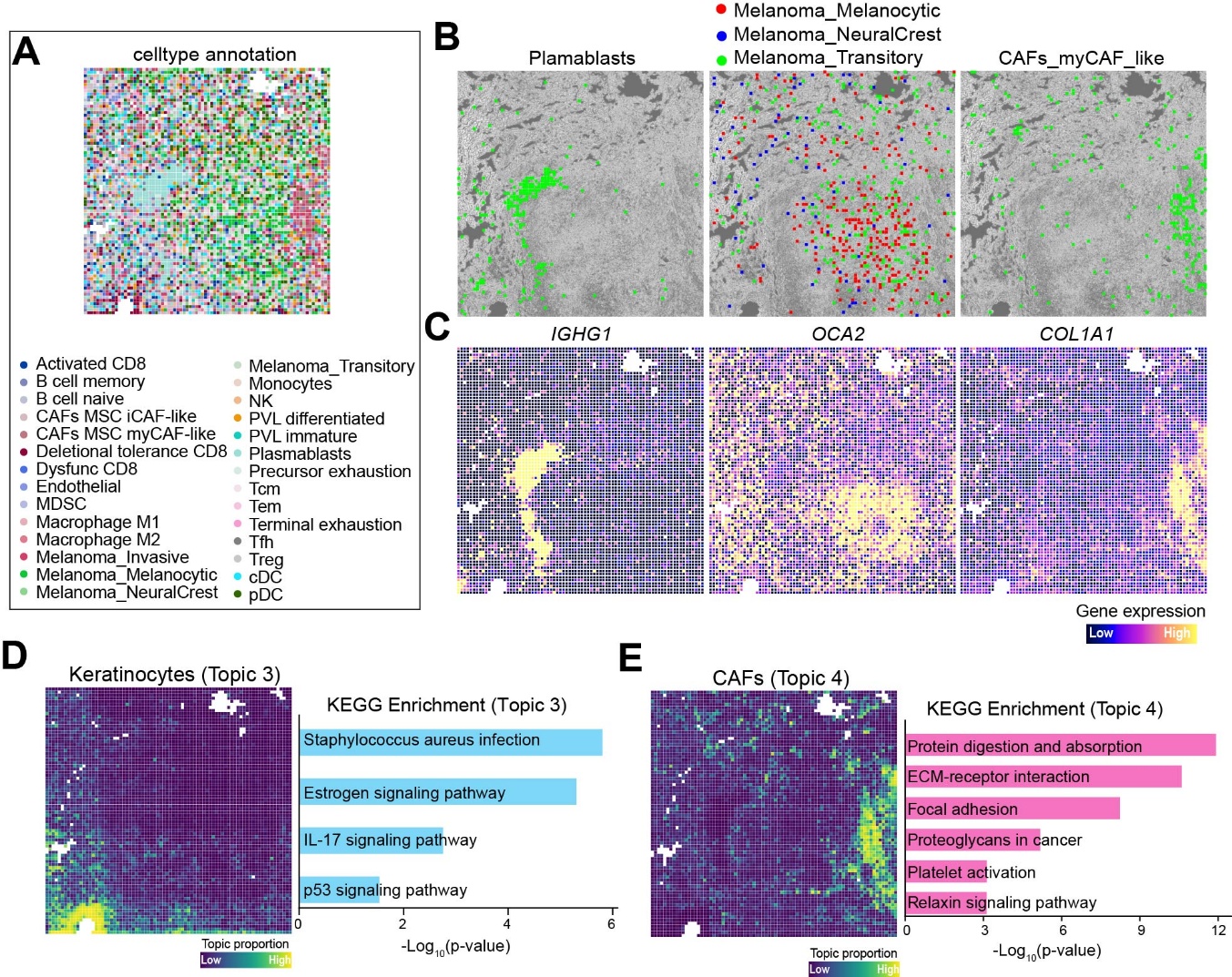

**Figure S6. Spatially resolved cell types and topic modeling in melanoma tissue**

(A) Spatial cell-type annotation derived from spatial transcriptomic profiling using reference-free annotation. Individual pixels are colored according to inferred cell identities.

(B) Spatial mapping of selected cell types, including plasmablasts (left), melanoma subtypes (melanocytic, neural crest–like, and transitory; middle), and myCAF-like cancer-associated fibroblasts (right).

(C) Spatial maps for *IGHG1*, *OCA2*, and *COL1A1*, representing markers of plasmablasts, melanocytic melanoma, and fibroblasts, respectively

(D) Topic modeling highlights a keratinocyte-enriched topic (Topic 3), shown as spatial topic proportion scores across the tissue (left) and KEGG pathway enrichment analysis (right).

(E) A fibroblast-associated topic (Topic 4) is shown by its spatial topic distribution (left) and corresponding KEGG pathway enrichment analysis (right).

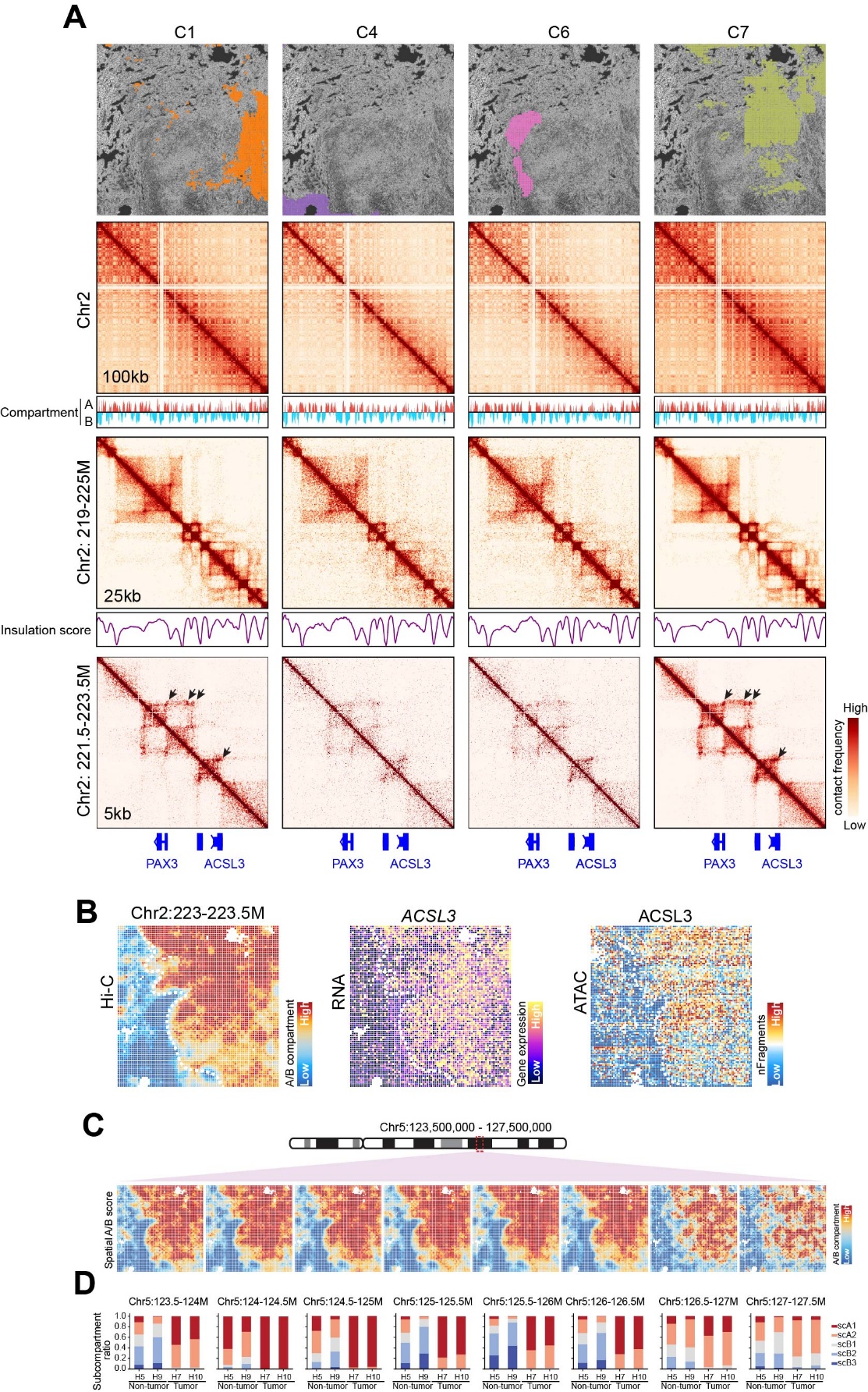

**Figure S7. Spatially resolved chromatin architecture and gene regulation at the *PAX3*–*ACSL3* locus in melanoma**

(A) Representative spatial domains (C1, C4, C8, C7) mapped onto tissue morphology (top). For each domain, multi-scale spatial Hi-C maps display chromatin interactions at 100, 25, and 5 kb resolution across the chr2: 21–23 Mb region encompassing *PAX3* and *ACSL3*. Below each map, compartment (A/B) profiles and insulation scores highlight local chromatin domain organization. High-resolution contact maps reveal variable loop formation (arrows) between *PAX3* and *ACSL3*, differing across spatial domains.

(B) Integration of chromatin conformation, spatial RNA-seq, and spatial ATAC-seq signals at the *ACSL3* locus. Spatial Hi-C contact enrichment (left), *ACSL3* transcript abundance (middle), and chromatin accessibility (right) show coordinated spatial regulation across melanoma regions.

(C) Spatial chromatin contact patterns along chr2:125.3–127.5 Mb using spatial A/B compartment scores, illustrating region-specific variability in 3D genome organization.

(D) Quantification of subcompartment states across the same genomic windows shown in (C). Barplots summarize the proportion of subcompartment assignments within selected spatial Hi-C clusters.

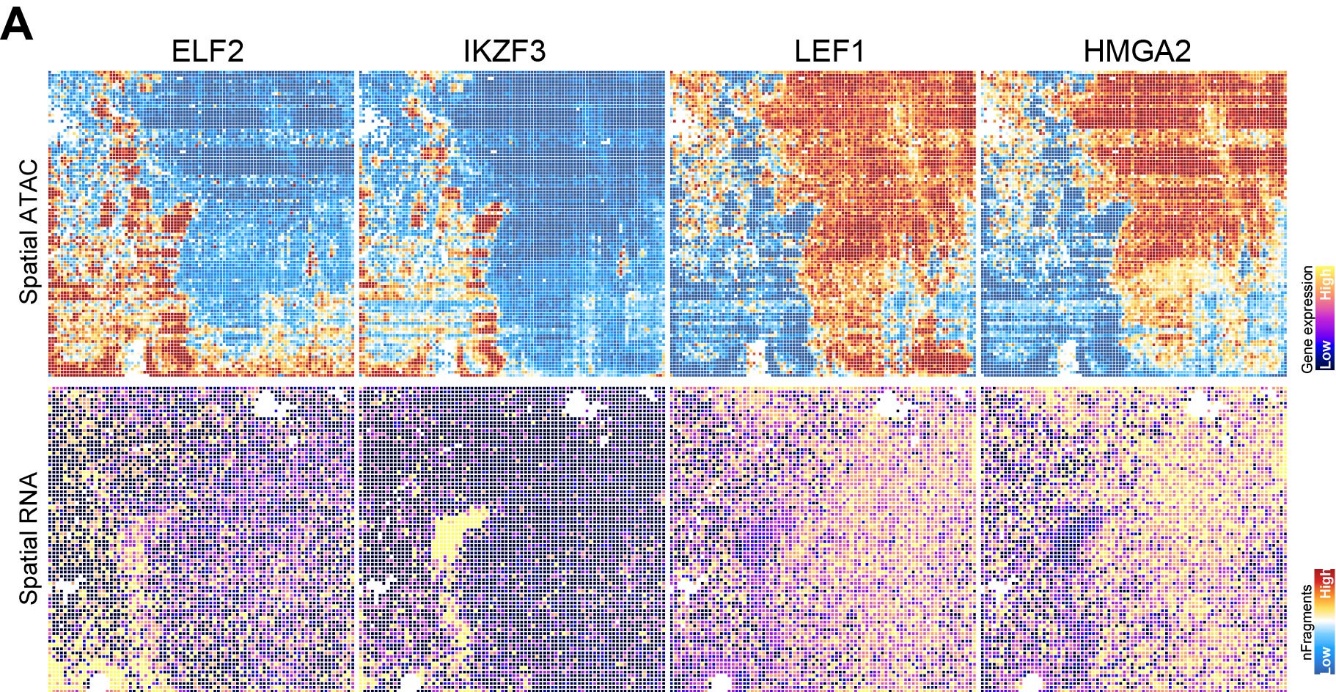

**Figure S8. Spatial distributions of chromatin accessibility and gene expression for selected regulators in melanoma**

(A) Spatial ATAC (top) and spatial RNA (bottom) patterns of selected regulators: ELF2 and IKZF3 for non-tumor regions, and LEF1 and HMGA2 for tumor regions.

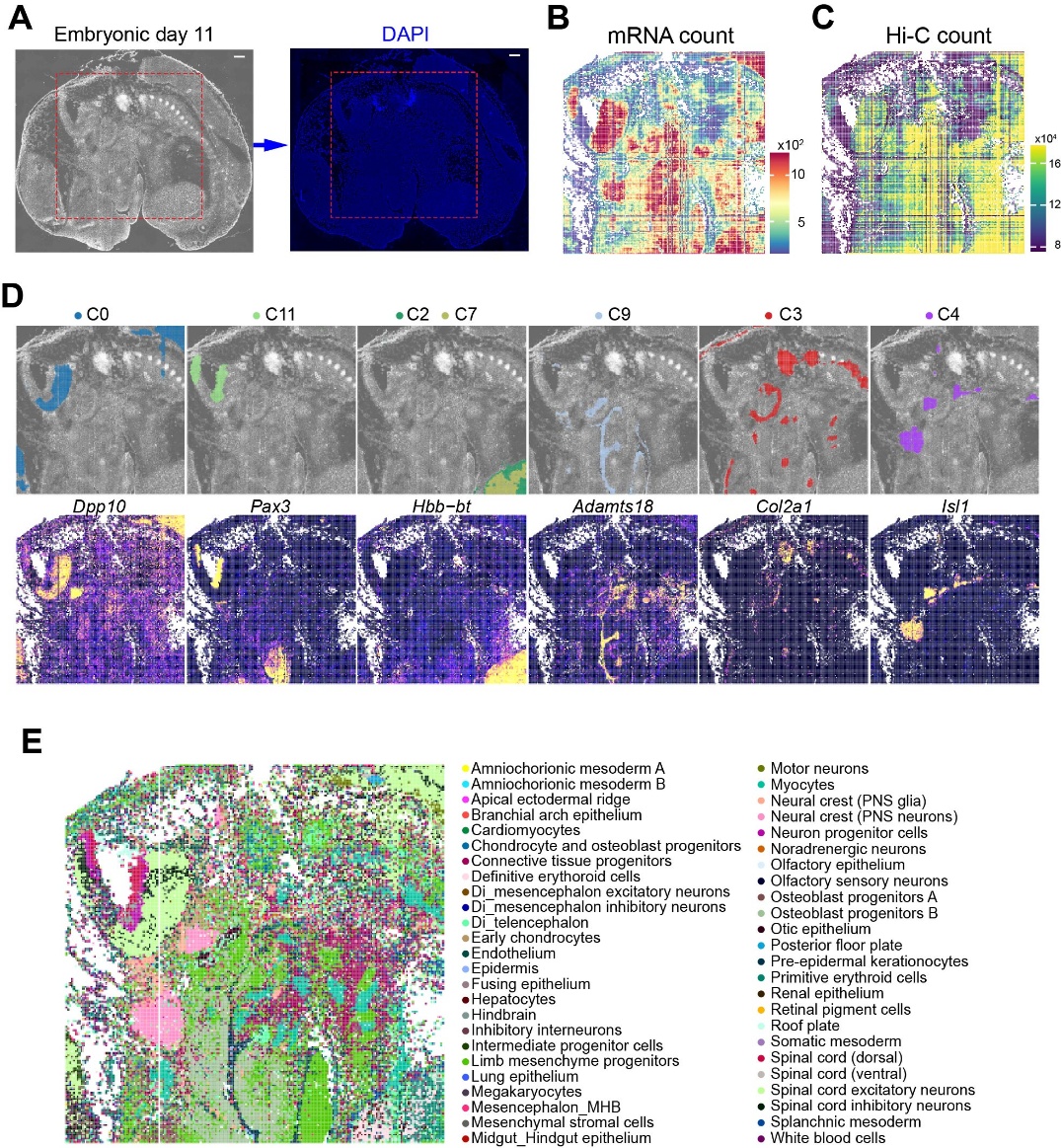

**Figure S9. Spatial multi-omic profiling of an E11.5 mouse embryo section using joint Hi-C–RNA mapping**

(A) Tissue overview and nuclear staining. Brightfield image of the E11.5 mouse embryo section prior to processing, with corresponding DAPI signal confirming intact tissue morphology and spatial registration. Scale bar: 500 μm.

(B-C) Spatial distribution of molecular signal intensities of spatial RNA (B) and Hi-C (C) count maps across the tissue section.

(D) Integrated spatial Hi-C/RNA domains with corresponding marker gene validation. Upper panel: spatial distribution of representative integrated clusters and lower panel: spatial expression patterns of selected marker genes—*Dpp10*, *Pax3*, *Hbb-bt*, *Adamts18*, *Col2a1*, and *Isl1*.

(E) Spatial cell-type annotation. Cell-type identities mapped from single-cell RNA-seq reference profiles onto each spatial pixel, revealing diverse embryonic lineages across the tissue section.

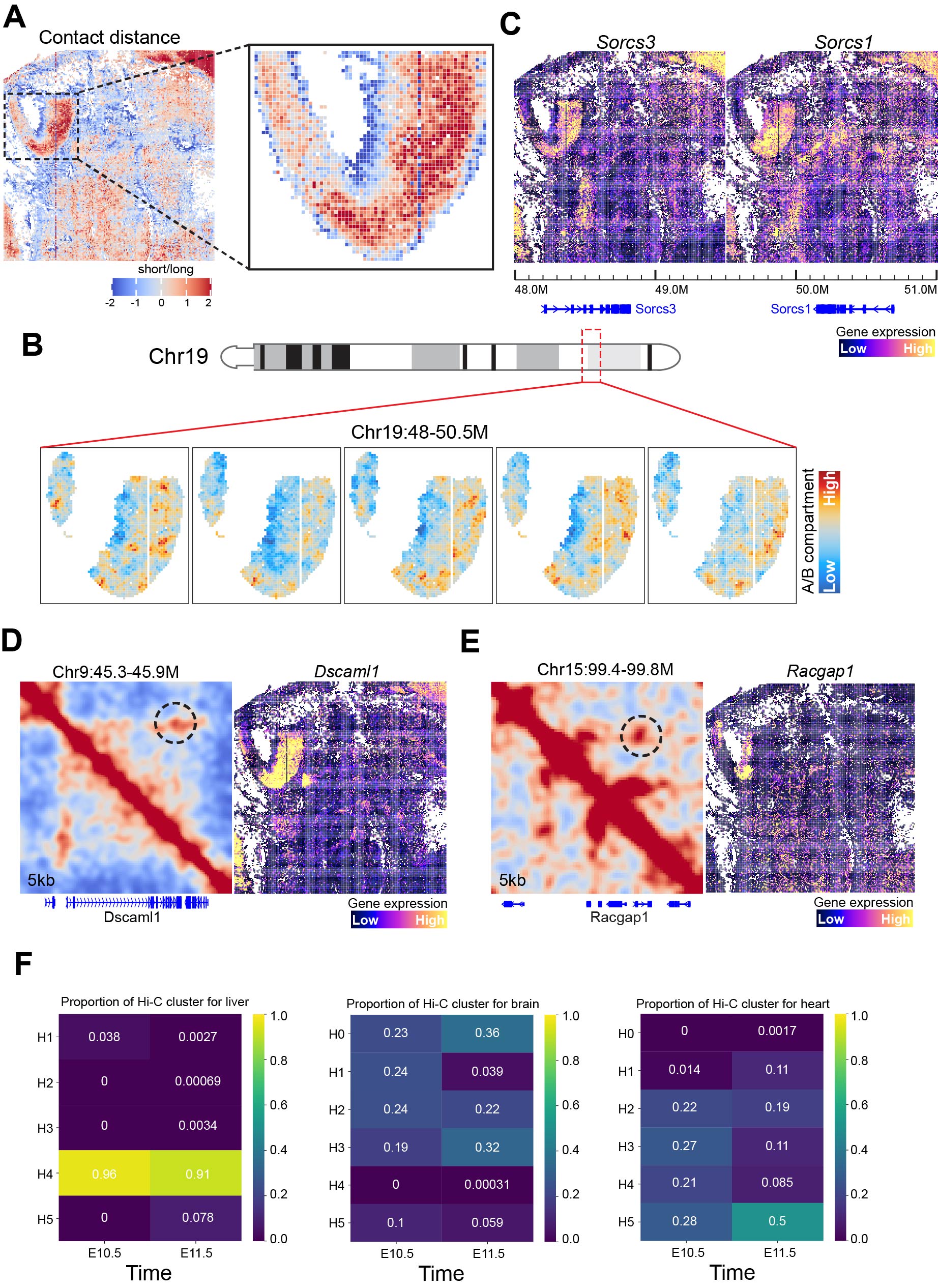

**Figure S10. Spatial chromatin remodeling and transcriptional changes during neural lineage progression and early organogenesis**

(A) Spatial variation in short- versus long-range chromatin contacts. Map of the cis short/long contact distance ratio across the ventral hindbrain, revealing a gradual increase along the neural progenitor–to–neuron differentiation trajectory. Zoomed inset highlights localized structural transitions associated with neuronal maturation.

(B) Spatial A/B compartment remodeling on chromosome 19. Top: schematic of chromosome 19 with the zoomed 48–50.5 Mb region indicated. Bottom: spatial A/B compartment scores across the embryo for subregions of Chr19, showing progressive acquisition of A-compartment features in differentiating neuronal populations.

(C) Gene expression of neural-associated loci. Spatial expression patterns of *Sorcs3* and *Sorcs1* across the same Chr19 region, demonstrating increased transcription in areas exhibiting A-compartment strengthening.

(D-E) Fine-scale chromatin interactions at lineage-specific genes.
High-resolution (5 kb) Hi-C maps show strengthened loops at the neuronal gene *Dscaml1* (D) and increased interaction intensity at the progenitor gene *Racgap1* (E), each matching their respective spatial expression domains.

(F) Organ-specific chromatin state dynamics inferred from transcriptomic data. Proportions of Hi-C cluster identities mapped onto Stereo-seq transcriptomes from liver, brain, and heart at E10.5 and E11.5, reflecting asynchronous chromatin reorganization across organs during early development.
